# Decoupling Phase Separation and Fibrillization Preserves Activity of Biomolecular Condensates

**DOI:** 10.1101/2025.03.18.643977

**Authors:** Tharun Selvam Mahendran, Anurag Singh, Sukanya Srinivasan, Christian M. Jennings, Christian Neureuter, Bhargavi H. Gindra, Sapun H. Parekh, Priya R. Banerjee

**Author notes:** Correspondence should be addressed to P.R.B.

## Abstract

Age-dependent transition of metastable, liquid-like protein condensates to amyloid fibrils is an emergent phenomenon of numerous neurodegeneration-linked protein systems. A key question is whether the thermodynamic driving forces underlying reversible phase separation and maturation to irreversible amyloids are distinct and separable. Here, we address this question using an engineered version of the microtubule-associated protein Tau, which forms biochemically active condensates. Liquid-like Tau condensates exhibit rapid aging to amyloid fibrils under quiescent, cofactor-free conditions. In particular, the Tau condensate interface promotes fibril nucleation, thereby impairing condensate activity in recruiting tubulin and catalyzing microtubule assembly. Remarkably, a small molecule metabolite, L-arginine, selectively impedes condensate-to-fibril transition without perturbing phase separation in a valence and chemistry-specific manner. By enhancing condensate viscoelasticity, L-arginine counteracts age-dependent decline in the biochemical activity of Tau condensates. These results provide a proof-of-principle demonstration that small molecule metabolites can reinforce the metastability of protein condensates against a liquid-to-amyloid transition, thereby preserving condensate function.

## Introduction

Protein condensation via phase separation is facilitated by an interplay between chain solvation and multivalent sequence-encoded inter-molecular interactions^1, 2, 3, 4, 5, 6^. Typically, protein condensates possess physical properties akin to viscoelastic fluids^7, 8, 9^. In certain protein systems associated with neurodegenerative disorders (ND), including FUS, hnRNPA1, TDP-43, and Tau, phase separation is linked to age-dependent alterations in condensate dynamics, morphology, and structure—a process known as physical aging or maturation^10, 11, 12, 13, 14, 15, 16^. The aging of protein condensates is an emergent property of the system described in recent literature using the framework of a glass transition^8^, dynamical arrest of the dense phase^9^, and a fluid-to-amyloid fibril transition^17^. Importantly, several clinically relevant mutations in ND-linked proteins were found to accelerate the liquid-to-fibril transition of condensates^11, 12, 16, 18, 19^, indicating that protein phase separation may contribute significantly to disease-associated fibrillization^20, 21, 22, 23^. However, an open question remains: Are the thermodynamic driving forces governing protein phase separation and fibrillization related or distinct?

Emerging experimental and computational results suggest that protein condensates are metastable relative to a globally stable solid state^9, 17, 24^. This may explain why stress granules hosting multiple aggregation-prone proteins become irreversible with prolonged persistence^11, 25^. It is plausible that active cellular processes and molecular chaperones, previously shown to regulate protein phase behavior^26, 27, 28, 29, 30, 31^, could counteract the age-dependent liquid-to-solid phase transitions of biomolecular condensates. Utilizing sequence engineering approaches for the prion-like low complexity domain of hnRNPA1, it was recently shown that the physical aging of protein condensates is sequence-encoded and partially distinct from the interactions that drive condensate formation^24^, leading to the postulate that distinct molecular grammars might exist for phase separation and physical aging. This indicates that macromolecules (such as proteins, RNAs, etc.) and/or small molecules that can selectively modulate sequence-specific interactions underlying amyloid formation could, in principle, buffer condensate liquid-to-solid transition independent of protein phase separation (**Fig. 1a**). Chemical targeting via small molecules has already been shown to modulate protein condensate formation and disassembly^26, 32, 33, 34^. With the growing interest in targeting aberrant biomolecular condensates with small molecules^35, 36, 37, 38, 39^, a key focus is on using small-molecule approaches to disentangle the molecular driving forces of phase separation from the physical aging of condensates associated with ND-linked proteins.

**Figure 1.**
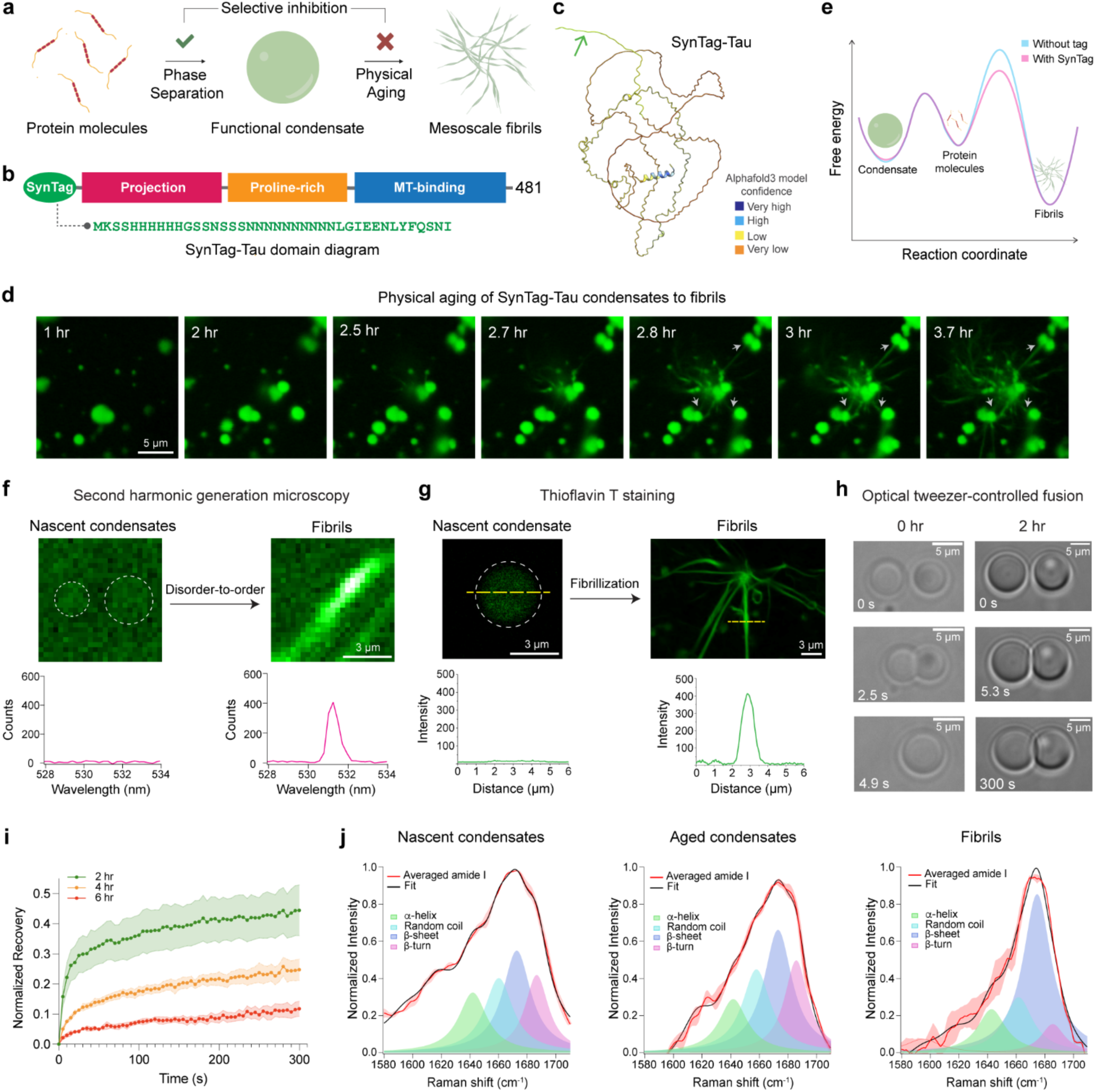
Engineered SynTag-Tau with a prionogenic tag undergoes phase separation coupled to fibril formation under cofactor-free quiescent conditions. **(a)** Schematic depicting the formation of phase-separated protein condensates and their time-dependent transition to form mesoscale amyloid fibrils. **(b)** Domain diagram of SynTag-Tau consisting of full-length Tau (2N4R isoform) with an N-terminal synthetic prionogenic tag (sequence provided in green). **(c)** Alphafold3^68^ predicted structure of SynTag-Tau. The green arrow marks the location of the tag. **(d)** Time-lapse imaging of Atto488-labeled SynTag-Tau condensates shows time-dependent morphological changes of the condensates to mesoscale fibrils. Grey arrows indicate the attachment of fibrils emerging from one condensate to neighboring condensates. Also, see **Supplementary Video 1**. **(e)** Schematic free energy landscape illustrating the relative stability of monomeric Tau, liquid-like condensates, and amyloid fibrils, with or without the prionogenic N-terminal SynTag. **(f)** Second harmonic generation (SHG) microscopy shows the absence of molecular ordering in nascent SynTag-Tau condensates (indicated by dashed circles; sample age = 2 hours) and the presence of spatial ordering in fibrils (sample age = 10 hours). The corresponding quantification is also shown. **(g)** Representative Thioflavin T (ThT) fluorescence line profiles from a nascent SynTag-Tau condensate (indicated by a dashed circle; sample age = 2 hours) and fibrils (sample age = 10 hours). **(h)** Optical tweezer-controlled fusion of nascent SynTag-Tau condensates (sample age = 30 mins, indicated as the 0-hour time point) and condensates at 2 hours of age. **(i)** FRAP measurements of SynTag-Tau condensates at various time points as indicated. The individual dots represent the mean value at each time point, based on 4-5 condensates from three independent replicates. Intensity error bars were plotted based on the standard error of the mean (S.E.M.) at each time point and are represented as the shaded regions. **(j)** Peak fitting of averaged amide I spectra obtained from broadband coherent anti-Stokes Raman (BCARS) hyperspectral imaging shows the structural profile of SynTag-Tau in condensates at 2 hours of age (nascent condensates) and at 6 hours of age (aged condensates), as well as fibrils formed at 10 hours of age. For panel (d), the concentration of SynTag-Tau used is 24 μM with 10% PEG 8000. For panels (f), (g), and (j), the concentration of SynTag-Tau used is 12 μM with 10% PEG8000. For panels (h) and (i), the concentration of SynTag-Tau used is 12 μM with 7.5% PEG8000. The buffer composition in all samples here is 10 mM HEPES (pH 7.4), 50 mM NaCl, 0.1 mM EDTA, and 2 mM DTT. Wherever applicable, the concentration of Atto488-labeled SynTag-Tau is 250 nM, and the concentration of ThT is 50 μM. Each experiment was independently repeated at least three times.

Motivated by these open questions, we sought to establish a model protein condensate system guided by the following criteria: (i) the ability to form fluid-like condensates that robustly age to globally stable solids under quiescent conditions, and (ii) condensates possess a tractable biochemical activity, enabling direct assessment of how physical aging affects condensate function. Establishing this benchmark will help differentiate between selectively disrupting condensate aging and broadly perturbing condensate formation, which could impact their biochemical activity. ND-linked proteins such as α-synuclein, FUS, and hnRNPA1 form condensates that undergo aging to fibrillar solids^12, 15, 17, 40^; however, the biochemical activities of condensates formed by these protein systems have not been established yet. Conversely, the microtubule-associated protein Tau, associated with a class of NDs called Tauopathies^41, 42, 43, 44, 45^, is known to form biochemically active condensates capable of microtubule (MT) assembly and stabilization implicated in intracellular trafficking in post-mitotic neurons^46, 47^. Importantly, multiple studies have hypothesized a link between the pathological cascade of Tau and Tau condensation, as pathogenic mutations have been shown to accelerate Tau condensate maturation^48, 49, 50, 51, 52^. These outstanding features provide a case for Tau as our model protein system. However, the aging of wild-type Tau condensates to amyloid fibrils is a kinetically sluggish process (**Supplementary Fig. 1**), hallmarked by the incredibly long timescale required for Tau aggregation and prion-like spread in vitro and in vivo^53, 54^. Tau is also well known for being highly soluble and recalcitrant to aggregation in vitro, often attributed to its hydrophilic nature^44, 55^. To enable Tau fibrillization within an experimentally accessible timescale, previous studies have used physicochemical perturbations, including negatively charged cofactors like heparin, shear stress, and modifications to the native protein, such as truncations or phosphorylation^49, 51, 56, 57^. These strategies impose unintended effects on Tau condensation and their activity in MT assembly^46, 58^, independent of condensate aging.

Here, we use SynTag-Tau, which comprises the full-length Tau protein (2N4R isoform) and a 40-amino-acid prionogenic N-terminal tag. Serendipitously, we found that this synthetic prion-like low-complexity sequence substantially reduces the Tau condensate-to-fibril nucleation barrier. SynTag-Tau undergoes molecular crowding-dependent phase separation in vitro to form biochemically active liquid-like condensates, similar to the untagged wild-type Tau protein^49^. Importantly, unlike the untagged counterpart, SynTag-Tau condensates undergo physical aging to mesoscale amyloid fibrils in the timescale of hours under quiescent conditions and in the absence of polyanionic cofactors, shear stress, or perturbations to the core microtubule-binding domain. Using the SynTag-Tau system, we uncovered that conditions that promote protein phase separation commensurately accelerate condensate aging to amyloid fibrils. SynTag-Tau condensate interface nucleates fibril formation, yielding a disorder-to-order transition with gradual changes in the condensate’s dynamical and structural properties. These age-dependent transitions in SynTag-Tau condensate physical properties accompany a progressive loss of biochemical activity in tubulin recruitment and catalyzing MT assembly. Remarkably, phase separation and physical aging in this system are decoupled by a naturally occurring small molecule metabolite abundantly present in the intracellular milieu, L-arginine (L-Arg). Our experiments show that the partitioning of L-Arg into the SynTag-Tau condensates prevents age-dependent changes in the condensate microenvironment, leading to an inhibition of fibril formation. Instead, it facilitates intra-condensate percolation transitions that enhance condensate metastability. L-Arg treatment of SynTag-Tau condensates preserves their activity in MT assembly. Finally, we show that L-Arg displays a similar activity to counteract the amyloid formation of untagged WT Tau condensates. Overall, this proof-of-principle study demonstrates that small molecule metabolites can stabilize protein condensates, preventing their transition from a liquid state to amyloid formation and thereby maintaining their function.

## Results and Discussion

### Physical aging of SynTag-Tau condensates leads to the formation of ordered amyloid fibrils

Wild-type (WT) Tau form condensates in physiologically relevant buffer conditions in vitro. These condensates are biochemically active^46^, but they do not transition to amyloid fibrils under quiescent conditions, thereby limiting our investigations (**Supplementary Fig. 1a, b**). Recent reports employing a protein sequence engineering strategy have enabled probing the interplay between phase separation and amyloid formation on an experimentally accessible timescale and resolution that is otherwise difficult to probe^59, 60^. These approaches exploit the principle that in many amyloid-forming protein systems, fibril formation is constrained by high nucleation barriers, which dictate the probability and kinetics of self-assembly^61^. Inspired by this phenomenon, we sought to design a sequence-engineered variant of Tau that selectively lowers the nucleation barrier for amyloid formation while preserving its intrinsic phase separation behavior and MT assembly function. Previous reports indicate that the fibrillar self-assembly of amyloidogenic proteins can be reinforced by fusing to a compact prionogenic low-complexity sequence^62, 63, 64^. Fusion to prion-like sequences flanking an amyloid-forming protein can act as soluble entropic bristles that favor the prion-like spread of protein aggregates without directly contributing to the primary nucleation mediated by the amyloid core^65^. In the current study, we adapted the strategy of adding a prion-like low-complexity sequence as an N-terminal tag to the Tau protein (**Fig. 1b**). We synthesized SynTag-Tau, which consists of the full-length Tau protein (2N4R isoform) with an amino-terminal tag containing a synthetic prionogenic sequence of 40 amino acid residues^66, 67^ (**Fig. 1b; Supplementary Fig. 2; Supplementary Table 1**). The design of this specific tag is serendipitous. Alphafold3^68^ predicted structure of SynTag-Tau indicates that this prion-like tag and most of the protein are intrinsically disordered, similar to that of the untagged Tau protein (**Fig. 1c; Supplementary Fig. 3**).

We first set out to characterize the formation and maturation of SynTag-Tau condensates utilizing a previously reported protocol of in vitro reconstitution in 10 mM HEPES (pH 7.4), 50 mM NaCl, 0.1 mM EDTA, 2 mM DTT, and a molecular crowder (10% PEG8000)^49, 52^. We tracked age-dependent morphological changes of SynTag-Tau condensates, doped with Atto488 (A488)-conjugated SynTag-Tau, by laser scanning confocal fluorescence microscopy (**Fig. 1d**). Within 2 to 3 hours (hr) after sample preparation, we observed fibrils from SynTag-Tau condensates growing in a heterogeneous manner*—*visible fibrils originated from some condensates, while others did not at this early time point. The fibrils nucleated from one condensate grow into the dilute phase and attach to neighboring condensates, inducing further nucleation and propagation of fibrillar assemblies (**Fig. 1d; Supplementary Video 1**). This observation appears to be consistent with a recent fibril growth model in the presence of condensates^24, 69^, where the growth of fibrils occurs in the dilute phase. Such a process engenders an efflux-generated gradient of protein molecules from nearby condensates, thereby creating a network of fibrils with condensates serving as nodes. Shuffling the primary sequence of the prionogenic tag without altering the overall sequence composition in a way that lowers the PLAAC score abrogates condensate aging to fibrils (**Supplementary Fig. 4**). Truncation of the tag sequence to limit it to the portion with the highest weighting in PLAAC score also renders the protein unable to trigger condensate-to-fibril transition (**Supplementary Fig. 5**). Thus, the specific composition and patterning of this synthetic prion-like tag sequence appear to be critical for lowering the nucleation barrier for the condensate-to-fibril transition (**Fig. 1e**). As a further validation, we sought to test whether the SynTag could serve as the primary driver of SynTag-Tau fibril formation. ZipperDB^70^ analysis revealed that in comparison to the native Tau sequence, the putative steric zippers in SynTag are outweighed in absolute score (Rosetta energy expressed as kcal/mol) and frequency, i.e., number of steric zippers (**Supplementary Fig. 6**). This suggests that the primary steric zippers capable of driving fibril nucleation likely originate from the native Tau sequence. However, without structural studies, we cannot rule out the possibility that the putative zipper motifs in SynTag act as auxiliary steric zippers that facilitate amyloid fibril formation.

We next employed in situ second harmonic generation (SHG), a non-linear label-free optical phenomenon ideal for probing non-centrosymmetric structures with molecular ordering^71^. We observed that SynTag-Tau fibrils, but not SynTag-Tau condensates, contain spatially ordered structures (**Fig. 1f**). This bears a resemblance to the SHG signal originating from bona fide amyloid fibrils observed in the brain of a transgenic model of Alzheimer’s^72^. To evaluate whether these ordered fibrils nucleated by condensates are indeed enriched in cross β-sheet structures, we employed an amyloid-sensitive dye, Thioflavin T (ThT). SynTag-Tau fibrils but not nascent SynTag-Tau condensates showed detectable ThT fluorescence (**Fig. 1g**). To probe the condensate-to-fibril transition in further detail, we measured the (a) timescale for droplet relaxation using a dual-trap optical tweezer, and (b) molecular mobility using fluorescence recovery after photobleaching (FRAP). We observed that the SynTag-Tau condensates undergo an age-dependent dynamical arrest, hallmarked by progressively slower condensate fusion and slower FRAP recovery, prior to the formation of visible amyloid fibrils in the mesoscale (**Fig. 1h, i**). At the molecular level, we hypothesized that a disorder-to-order conformational change of SynTag-Tau is likely to aid in dynamical arrest and concomitant fibrillization. To obtain an age-dependent structural fingerprint of SynTag-Tau molecules in the dense phase, we employed in situ spatially resolved broadband coherent anti-Stokes Raman scattering (BCARS)^73^, which is a label-free hyperspectral Raman imaging technique. Intra-condensate BCARS measurements revealed a progressive increase in the β-sheet content with condensate age, ultimately leading to amyloid fibrils that are predominantly enriched in β-sheet conformers (**Fig. 1j; Supplementary Table 2**).

So far, we have established that SynTag-Tau condensates physically age from a fluid-like phase to fibrillar solids enriched in β-sheet structures. Next, we asked whether these two processes are correlated, meaning how perturbation of SynTag-Tau condensate formation impacts SynTag-Tau fibrillization. Previous literature suggests that electrostatic interactions play a dominant role in Tau phase separation^56, 74, 75^, which can be modulated by tuning the ionic strength of the buffer. We observed that increasing ionic strength diminishes SynTag-Tau phase separation (measured by condensate area fraction). Interestingly, the kinetics of SynTag-Tau fibrillization decrease in a correlated manner (**Supplementary Fig. 7a, b**). Further, we tested this interplay by titrating the molecular crowder concentration, wherein high [PEG8000] favors protein phase separation propensity via depletion forces^76^. Upon increasing the concentration of the crowder, we find that SynTag-Tau phase separation is enhanced, judged by an increase in condensate volume fraction, and concomitantly, the timescale for condensate-to-fibril transition is reduced (**Supplementary Fig. 7c**). Taken together, these observations suggest that promoting SynTag-Tau condensation enhances fibrillization. Although the insights gained so far seemingly indicate an interplay between phase separation and fibril formation, a key question is how condensate physical aging impacts the function of SynTag-Tau condensates in tubulin recruitment and MT assembly.

### Condensate physical aging impairs biochemical activity

We utilized an in vitro tubulin recruitment and MT assembly assay^46^ (**Fig. 2a**) to quantify the impact of SynTag-Tau condensate aging on biochemical activity. We first tested the partitioning of tubulin alone in the absence of GTP in nascent and aged SynTag-Tau condensates. We find that the nascent condensates can recruit tubulin (labeled with HiLyte647) with homogeneous distribution throughout dense phase (**Fig. 2b; Supplementary Fig. 8**). Contrastingly, tubulin partitioning was attenuated in aged SynTag-Tau condensates with tubulin molecules mostly concentrating close to the interface (**Fig. 2c; Supplementary Fig. 8**). Next, we tested the age-dependent activity of SynTag-Tau condensates using in vitro MT polymerization assay (experimental schematic in **Fig. 2a**). With increasing condensate age (time = Δ*t*), we observed a gradual reduction in MT polymerization capacity (**Fig. 2d, e; Supplementary Fig. 9, 10, 11**). By the 16-hour time point, the ability of SynTag-Tau condensates to drive MT polymerization ceases completely. Consistently, tubulin partitioning experiments conducted using similar assay conditions as the MT polymerization assay revealed increasingly heterogeneous tubulin partitioning at later time points, which correspond to loss of activity (**Supplementary Fig. 12; Fig. 2e**). The ineffective partitioning of tubulin to condensates at later ages suggests that the characteristic mesh size might progressively decline with physical aging^77, 78^. Nevertheless, it is worth pointing out that in the 8 hour time point, despite observing a noticeable drop in MT assembly, we do not observe a dramatic blockade of tubulin partitioning via fluorescence microscopy, which might indicate that aging-induced changes in material state, as well as perturbed tubulin binding by conformationally altered SynTag-Tau within aged condensates, might underscore condensate loss-of-activity. Overall, the loss of activity of SynTag-Tau condensates with age demonstrates that the condensate material properties affect tubulin recruitment and MT assembly. Similar attenuation of SynTag-Tau condensate activity was observed (**Supplementary Fig. 13**) when we increased the molecular crowder concentration from 5% to either 7.5% or 10%, which accelerates SynTag-Tau condensate-to-fibril transition (**Supplementary Fig. 7c**) and was previously shown to modulate condensate material properties^76^.

**Figure 2.**
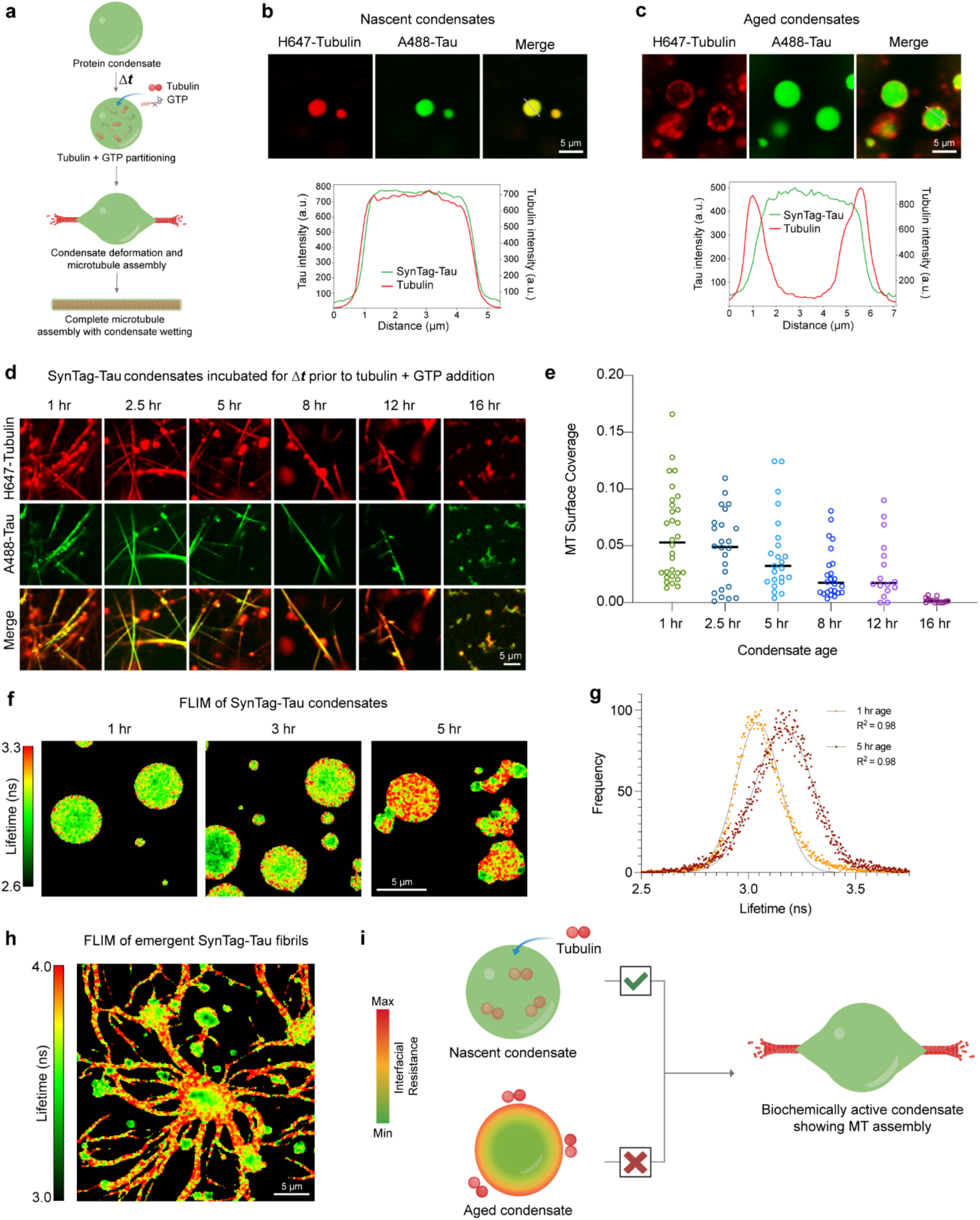
Age-dependent liquid-to-amyloid transition impairs SynTag-Tau condensate activity in microtubule assembly. **(a)** Schematic of tubulin recruitment and MT polymerization in Tau condensates. **(b)** Nascent SynTag-Tau condensates (age = 2 hours) in the presence of 50 nM HiLyte647-labeled tubulin show enrichment of tubulin in the dense phase. Corresponding line profiles are shown below. **(c)** Aged SynTag-Tau condensates (age = 4 hours) with the addition of 50 nM HiLyte647-labeled tubulin show altered tubulin partitioning to the condensate interface. Corresponding line profiles are shown below. **(d)** Condensate age-dependent MT polymerization assay in SynTag-Tau condensates. **(e)** MT surface coverage plot corresponding to panel (d). Horizontal lines at each time point represent the median value. The individual data points represent 14-30 measurements, based on three independent replicates. **(f)** Frequency-domain (FD) FLIM images of SynTag-Tau condensates at various time points highlight the age-dependent increase in the lifetime of Atto488-labeled SynTag-Tau molecules at the condensate interface, which eventually spreads to the condensate core. **(g)** Representative fluorescence lifetime distributions at 1 hour and 5 hour time points (since sample preparation). The points are the data, and the lines are Gaussian fittings with the corresponding goodness of fit (R^2^) indicated. **(h)** FD-FLIM map of aged SynTag-Tau condensates with emergent fibrillar assemblies (sample age = 8 hours). **(i)** Schematic of condensate physical aging-induced interfacial resistance in SynTag-Tau condensates, which perturbs partitioning of tubulin to the condensate dense phase and impairs MT assembly. The concentration of SynTag-Tau protein used in (b, c) is 24 μM with the following buffer composition, 10 mM HEPES (pH 7.4), 50 mM NaCl, 0.1 mM EDTA, and 2 mM DTT along with 10% PEG8000. In (f-h), the concentration of SynTag-Tau protein is 12 μM with the same buffer composition as (b, c). In (d), the concentration of SynTag-Tau used is 12 μM with the following buffer composition, 80 mM PIPES (pH 6.9), 2 mM MgCl_2_, 0.5 mM EGTA, and 2 mM DTT along with 5% PEG8000 crowder. In (d), the tubulin concentration is 5 μM, and the concentration of GTP is 1 mM. Wherever applicable, the concentrations of Atto488-labeled SynTag-Tau and HiLyte647-labeled tubulin are 250 nM and 500 nM (unless specified otherwise), respectively. Each of these experiments was independently repeated at least three times.

Recent reports suggest that the interface of protein condensates plays a critical role in nucleating the liquid-to-amyloid transition^17, 79, 80^. This is presumed to stem from the unique conformational ensemble of the protein molecules at the condensate interface, in contrast to the dilute or dense phase^81^. Therefore, we asked whether SynTag-Tau condensate interfaces play a similar role in fibril formation. To address this, we employed frequency-domain fluorescence lifetime imaging (FLIM) with Atto488-labeled SynTag-Tau. FLIM is a highly sensitive imaging technique that reports on the spatial heterogeneity in the molecular microenvironment of a fluorescent probe. In our experiment, we measured the lifetime of the Atto488-labeled SynTag-Tau to interrogate the local environment of protein molecules within the condensate as they undergo age-dependent morphological transitions. We find that there is a clear lifetime signature of SynTag-Tau molecules on the interface of the condensates at an early age, which strikingly spreads to the condensate core as the age-dependent transformation of liquid protein condensates to amyloid fibrils takes place (**Fig. 2f, g**). The lifetimes of SynTag-Tau molecules at the condensate interface are longer than those of the bulk, closer to the lifetimes of the fibrils (**Fig. 2h**). These data suggest that SynTag-Tau condensate interfaces nucleate amyloid formation. We infer that the altered tubulin partitioning in nascent versus aged protein condensates (**Fig. 2b, c; Supplementary Fig. 12**) is due to the physical aging of SynTag-Tau condensates that begin at the interface (**Fig. 2i**). Disabling and/or delaying the fluid-to-fibril transition of protein condensates could potentially protect against condensate loss-of-activity.

### Naturally occurring small molecule metabolites modulate Tau condensate physical aging

Identifying small molecule inhibitors of protein fibrillization has been an active area of research for over two decades. In the context of phase separation, small molecules, such as 4,4’-dianilino-1,1’-binaphthyl-5,5’-disulfonic acid (bis-ANS) and adenosine triphosphate (ATP), were shown recently to modulate the phase behavior of ND-linked proteins^26, 32^. These studies and mechanistic perspectives gleaned by quantitative frameworks such as polyphasic linkage^82^ and heterotypic buffering^83, 84^ have postulated that tuning condensate material properties may prevent their aberrant behavior. Motivated by this perspective, we set out to test whether SynTag-Tau condensate maturation timescale can be tuned independently of SynTag-Tau phase separation through small molecule treatment. We first tested generic chemical disruptors of protein-protein interactions and naturally occurring small molecule metabolites at low millimolar concentrations. Remarkably, we found that physiologically abundant cationic amino acids, L-arginine (L-Arg) and L-lysine (L-Lys), inhibit SynTag-Tau condensate-to-fibril transition without perturbing SynTag-Tau phase separation (**Fig. 3a**). The concentrations at which L-Arg showed such activity is similar to its physiologically relevant concentrations^85, 86^. In the presence of anionic amino acids such as L-glutamic acid (L-Glu) and L-aspartic acid (L-Asp), however, the appearance of SynTag-Tau fibrils seemed to accelerate. The observed tunability of SynTag-Tau liquid-to-fibril transition by these small molecule metabolites is likely owed to their charge and the side chain chemistry as L-proline (L-Pro), which cannot participate in electrostatic interactions, failed to show any modulatory effect (**Fig. 3a**). The inhibitory effect of L-Arg and L-Lys was also not recapitulated by small molecules with pleiotropic effects on protein-protein interactions, such as those that disrupt protein backbone hydrogen bonding interactions, including guanidinium hydrochloride (GnHCl) and urea (**Fig. 3a**). The chemical chaperone trimethylamine N-oxide (TMAO) previously reported to disfavor fibrillization of TDP-43^27^ was unable to prevent SynTag-Tau phase separation coupled fibrillization (**Fig. 3a**). Conversely, ATP appears to hinder fibril formation, however, the condensate size and morphology are substantially affected (**Fig. 3a**).

**Figure 3.**
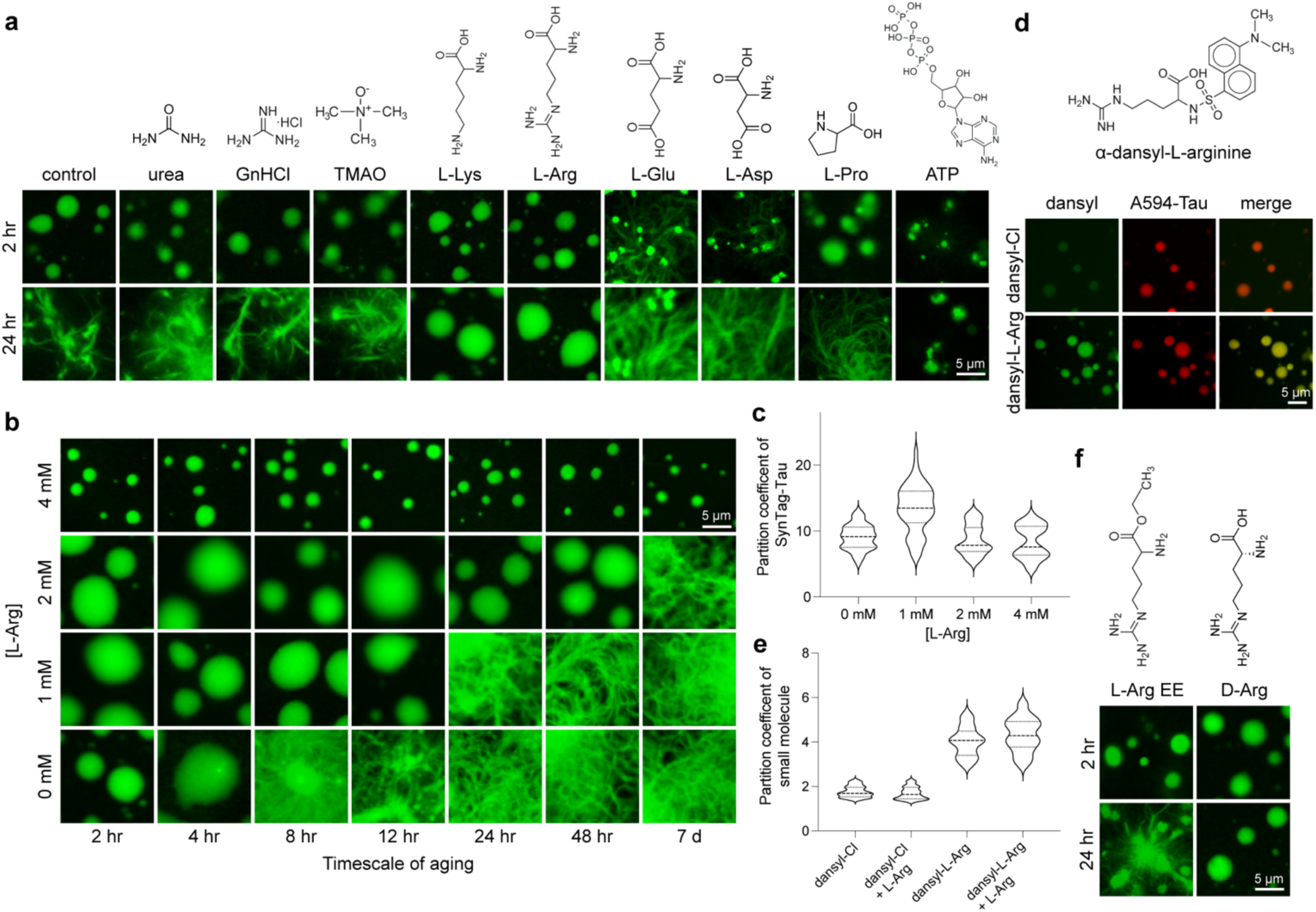
Small molecule metabolites can decouple phase separation and fibril formation in a chemistry-specific manner. **(a)** Effect of naturally occurring small molecule metabolites, chaotropic compounds, and small molecule modulators of protein-protein interactions on phase separation and fibrillization of SynTag-Tau. The chemical structure of the respective small molecule is also shown. **(b)** Dose-dependent effect of L-Arg in inhibition of SynTag-Tau condensate-to-amyloid transition. **(c)** The corresponding partition coefficient analysis from fluorescence images as shown in (b). The thick dashed line represents the median based on data of 53 condensates from three independent replicates, whereas the thinner dotted lines above and below represent the upper and lower quartiles, respectively. **(d)** (top) Chemical structure of α-dansyl-L-arginine (dansyl-L-Arg), a fluorescently labeled analog of L-Arg. (bottom) Partitioning of α-dansyl chloride (dansyl-Cl) or dansyl-L-Arg doped along with L-Arg into SynTag-Tau condensates, visualized using Alexa594- (A594) labeled SynTag-Tau, at 2 hours sample age. **(e)** Partition coefficient analysis of dansyl-Cl or dansyl-L-Arg, either with or without L-Arg to SynTag-Tau condensates. The thick dashed line represents the median based on data of 32 condensates from three independent replicates, whereas the thinner dotted lines above and below represent the upper and lower quartiles, respectively. **(f)** L-Arg ethyl ester (L-Arg EE), a derivative of L-Arg that lacks a carboxyl group, failed to prevent condensate aging to fibrils. D-arginine (D-Arg) treated condensates do not transition to fibrils, similar to the L-Arg condition shown in (a). The composition of these samples is 12 μM SynTag-Tau in buffer containing 10 mM HEPES (pH 7.4), 50 mM NaCl, 0.1 mM EDTA, and 2 mM DTT along with 7.5% PEG8000. Small molecules were introduced prior to the induction of protein phase separation, and their concentrations used in these experiments are 2 mM unless specified otherwise. Wherever applicable, the concentration of Atto488-/Alexa594-labeled SynTag-Tau is 250 nM, and the concentration of dansyl-L-Arg/dansyl-Cl is 500 nM (doped with/without L-Arg to make up a total small molecule concentration of 2 mM). Each of these experiments was independently repeated three times.

We next turned our attention to L-Arg to determine its dose-dependent efficacy in perturbing SynTag-Tau fibrillization (**Fig. 3b**). At the lowest L-Arg concentration tested (1 mM), the partition coefficient of Atto488-labeled SynTag-Tau molecules in the dense phase is higher compared to the untreated condition. However, increasing the L-Arg concentration to 2 mM or 4 mM did not dramatically alter the partition coefficient relative to the untreated condition (**Fig. 3b, c**). This data suggests that SynTag-Tau phase separation is not negatively impacted by 1 mM to 4 mM L-Arg. Notably, at 2 mM L-Arg concentration, SynTag-Tau condensate size and morphology looked similar to those of the untreated condition, but the timescale of fluid-to-fibril transition shifted from 8 hours to 7 days (**Fig. 3b, c**). At 4 mM L-Arg, however, we observed some differences in SynTag-Tau condensate size and surface wetting behavior compared to the untreated conditions (**Fig. 3b, c**) despite the SynTag-Tau partition coefficient being largely similar. In order to ascertain whether the reduced condensate sizes stem from either altered condensate coarsening dynamics, inhibition of SynTag-Tau phase separation or both, we conducted a quantitative analysis of condensate area fraction using confocal fluorescence microscopy (**Supplementary Fig. 14**). We found that L-Arg concentrations of 1 mM and 2 mM yielded condensate sizes that are not significantly different than that of the untreated condition (0 mM). However, in the case of 4 mM L-Arg treatment, we observed a significantly diminished condensate area fraction, indicating that L-Arg at high concentrations (above 2 mM) inhibits SynTag-Tau phase separation and potentially affects condensate coarsening. The selective inhibition of fibrillization without abrogating condensate formation reveals a distinction in the molecular driving forces underlying these two processes, indicating that phase separation and fibrillization are biochemically separable. This decoupling phenomenon is distinct from global perturbation to condensate formation, as seen with the effect of modulating ionic strength or molecular crowding (**Supplementary Fig. 7**).

Next, we probed the partitioning behavior of L-Arg into SynTag-Tau condensates utilizing α-dansyl-L-arginine (dansyl-L-Arg), a fluorescent analog of L-Arg (**Fig. 3d**). Using dual-color fluorescence microscopy with SynTag-Tau condensates visualized with Alexa594-conjugated protein, we find positive partitioning with a partition coefficient, k ∼ 4, of dansyl-L-Arg within SynTag-Tau condensates (**Fig. 3e**). This partition coefficient value is consistent with the known partition coefficients of similar small molecule metabolites in condensates^39^. The partitioning of dansyl-L-Arg is not due to the dansyl group as partitioning of α-dansyl chloride to SynTag-Tau condensates is much weaker (k ∼ 1.5) (**Fig. 3d, e**).

To investigate how the inhibitory effect of L-Arg on SynTag-Tau condensate aging is related to its chemical properties, we tested L-Arg ethyl ester (L-Arg EE) and observed no impact on SynTag-Tau condensate aging (**Fig. 3f**). Experiments with D-arginine (D-Arg) yielded similar results as L-Arg, revealing that the stereospecificity of arginine is not critical to its inhibitory activity (**Fig. 3f**). Next, we tested whether multivalent polymers of lysine or arginine as well as charged polyamines such as spermine and spermidine could impose an enhanced inhibitory potential to condensate-to-fibril transformation relative to small molecules (L-Arg/L-Lys). Intriguingly, treating condensates with multivalent cationic molecules did not offer any improvement in fibrillization timescale relative to the small molecule treatment (**Supplementary Fig. 15; Fig. 3a**). Given the observation that the positive hits in the small molecule screen are basic amino acids with chemical moieties of high pKa (>7), L-Lys (pKa of ε-amino group = 10.5), and L-Arg (pKa of guanidino group = 12.4), we wondered at the possibility that high pKa could be an important chemical property that confers protection against condensate-to-fibril transition. However, the remaining small molecules with basic moieties failed to achieve a similar effect. For example, guanidinium hydrochloride (pKa of guanidinium group = 12.6), L-arginine ethyl ester (pKa of guanidino group = 12.4), and spermine and spermidine (pKa across multiple amine groups = ∼10.7). Despite sharing highly basic, i.e., high pKa chemical moieties, several of these compounds failed to inhibit SynTag-Tau fibrillization selectively (**Fig. 3a**; **Supplementary Fig. 15**), which suggests that strongly basic (high pKa) functional groups alone are a poor predictor of inhibitory potential. Therefore, we conclude that the observed inhibitory activity of L-Arg, close to its physiological concentration, is chemistry-, valence-, and charge-specific, but does not depend on the chirality.

### L-Arg prevents cross β-sheet formation and nucleation of fibrils at the condensate interface

Given that L-Arg can prevent age-dependent changes in SynTag-Tau condensate morphology (**Fig. 3**), we next sought to probe its effect on the structure and dynamics of SynTag-Tau condensates. Using the ThT fluorescence assay as a function of time, we observed that the age-dependent increase in the signal originating from amyloid fibrils is notably less pronounced for condensates in the presence of L-Arg (**Fig. 4a**). This observation suggests that L-Arg inhibits conformational conversion of disordered SynTag-Tau molecules within condensates to cross-β-rich conformers, preventing nucleation of amyloid fibrils. One caveat of imaging condensates at a single plane using ThT fluorescence is the lack of spatial information in the z-axis that would be critical to resolve whether fibrils are indeed nucleated at condensate interfaces or if they localize to condensate interiors. We tested this by capturing z-slices, utilizing A594-labeled SynTag-Tau and ThT fluorescence, in samples of untreated SynTag-Tau condensates. We find that ThT fluorescence is preferentially enriched at condensate interfaces with amyloid fibrils growing into the dilute phase, but not condensate interiors (**Fig. 4b**). At a relatively higher z position (0.95 μm), i.e., approaching the topmost interface of condensates (with respect to the coverslip surface), the ThT signal originating from fibrils becomes more prominent, in line with interfacial nucleation of fibrils. This is consistent with the recently proposed framework that condensate interiors could act as sinks of amyloidogenic proteins; however, their interfaces present the risk of fibril nucleation, with fibril growth occurring outward into the dilute phase^24^. On the other hand, L-Arg-treated SynTag-Tau condensates fail to show any notable ThT fluorescence across z-sections (**Fig. 4c**). For further examination, we employed BCARS hyperspectral imaging and observed that the amide I band of SynTag-Tau in nascent condensates is fairly similar in either absence or presence of L-Arg (**Fig. 4d, e; Supplementary Table 2**). However, the amide I band of aged SynTag-Tau condensates treated with 2 mM L-Arg appears distinct from that of untreated condensates at the same age. The most dramatic difference is the decrement in the mean population of β-sheet conformers in L-Arg-treated condensates (39.3%) versus the untreated condition (47.2%). We also note a moderate increase in non-β conformations in small molecule-treated condensates relative to untreated condensates, namely random coil (37.4% vs. 32.9%, respectively) and α-helix (23.3% vs. 19.9%, respectively) (**Fig. 4e**). This may suggest an exciting possibility that L-Arg might slow down the SynTag-Tau condensate liquid-to-fibril transition by populating alternative ensemble conformational states of SynTag-Tau. Follow-up work is necessary to explore this possibility further.

**Figure 4.**
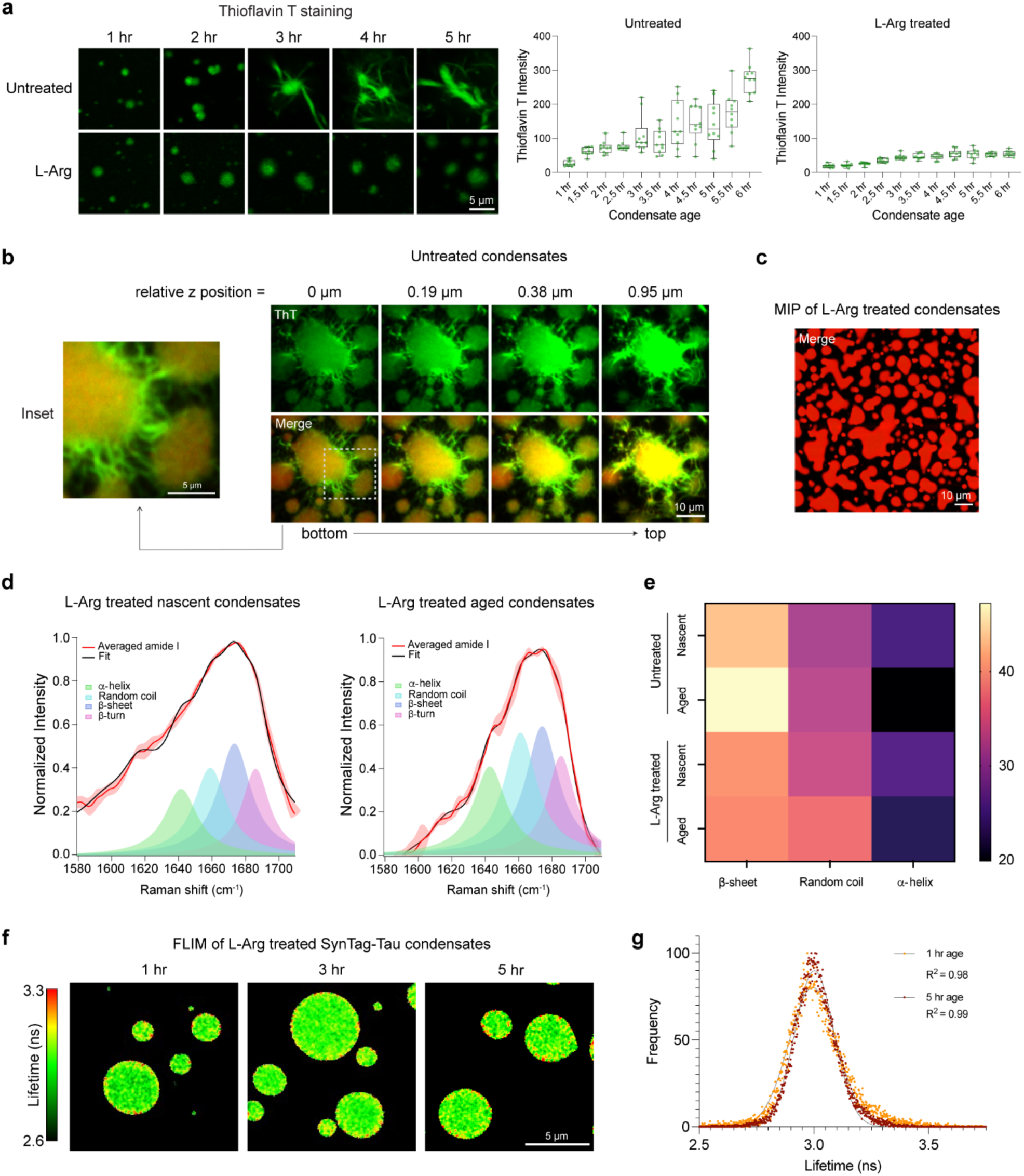
L-Arg prevents cross β-sheet formation and nucleation of fibrils at the condensate interface. **(a)** ThT fluorescence of condensates at different sample ages without (untreated) and with 2 mM L-Arg. Corresponding plots of ThT fluorescence intensities are shown on the right. The contrast was adjusted independently across these images to better visualize condensates that are very dim. The center line represents the median, and the individual data points represent measurements from 10 condensates across three independent replicates. **(b)** Individual Z-slices of untreated SynTag-Tau condensates visualized using ThT (green) as well as A594-labeled SynTag-Tau (red). The indicated z-position is relative to the leftmost z-slice in the panel. The inset shown represents the area demarcated by the white-dashed box. Sample age is 48 hours. **(c)** Maximum intensity projection (MIP) image of 2 mM L-Arg treated SynTag-Tau condensates with A594-labeled SynTag-Tau and ThT. Sample age is 48 hours. Notably, the ThT fluorescence intensity is substantially lower relative to the A594-labeled SynTag-Tau fluorescence for these condensates, in contrast to that of the untreated SynTag-Tau condensates, suggesting a lack of fibrils with L-Arg treatment. **(d)** Peak fitting of averaged amide I spectra obtained from BCARS hyperspectral imaging of L-Arg-treated SynTag-Tau condensates reveals changes in protein molecular conformations from 2 hours of age (nascent condensates) to 6 hours of age (aged condensates). **(e)** Heat map based on the amide I spectra obtained through BCARS hyperspectral imaging showing the mean percentages of protein conformations in untreated versus 2 mM L-Arg treated SynTag-Tau condensates at 2 hours (nascent) and 6 hours (aged) of age. The mean percentages based on measurements from three independent replicates represent the quotient of each conformation’s peak fitted area divided by the cumulative area of β-sheet, random coil, and α-helix. **(f)** FD-FLIM map of L-Arg treated condensates at various time points (since sample preparation). **(g)** Representative fluorescence lifetime distributions at 1 hour and 5 hour time points (since sample preparation). The points are the data, and the lines are Gaussian fittings with the corresponding goodness of fit (R^2^) indicated. The composition of these samples is 12 μM SynTag-Tau in buffer containing 10 mM HEPES (pH 7.4), 50 mM NaCl, 0.1 mM EDTA, and 2 mM DTT along with 10% PEG8000 crowder. The L-Arg concentration used in these experiments is 2 mM. Wherever applicable, the concentration of A488/A594 labeled SynTag-Tau is 250 nM, and the concentration of ThT is 50 μM. Each of these experiments was independently repeated three times.

To probe the spatially resolved SynTag-Tau condensate microenvironment with age in the presence of L-Arg, we performed FLIM measurements using an identical approach as outlined in **Fig. 2f**. When comparing with the SynTag-Tau condensates without L-Arg, we made two key observations. The first is that the lifetime of fluorescently labeled SynTag-Tau molecules does not change over the same timescale, and the interface of these condensates appears similar in terms of the fluorescence lifetime with respect to the SynTag-Tau molecules at the condensate core (**Fig. 4f, g**). These data collectively suggest that the mesoscale changes in the SynTag-Tau condensate morphology with age are inhibited by L-Arg. Finally, we tested the efficacy of L-Arg in the heparin-induced maturation of WT Tau condensates into amyloid fibrils. L-Arg proved effective in this in vitro model of condensate aging as well, indicating that the selective inhibitory potential of L-Arg against condensate-to-fibril transition does not stem from interactions between L-Arg and the synthetic prionogenic tag in SynTag-Tau but potentially via frustration of the interactions mediated by the zipper motifs in Tau (**Supplementary Fig. 16**). Contrarily, treatment with L-Glu resulted in progressively higher ThT fluorescence relative to the untreated condition, consistent with observations in the SynTag-Tau model system (**Supplementary Fig. 16**).

### Small molecule-induced enhancement in condensate viscoelasticity weakens fibrillization propensity

The effect of L-Arg in regulating condensate-to-fibril transition may stem from modulating the material properties of Tau condensates. To test this idea, we employed video particle tracking (VPT) nanorheology using 200 nm-sized fluorescent probe particles passively embedded inside WT Tau condensates (**Fig. 5a**). From the mean squared displacements (MSDs) of the fluorescent probes, we computed the frequency-dependent viscoelastic moduli of Tau condensates. The timescale of the dominantly viscous versus dominantly elastic behavior of Tau condensates can be deduced by estimating the crossover frequency from the frequency-dependent viscoelastic moduli. In the absence of L-Arg, Tau condensates are highly viscoelastic, with probe particles exhibiting sub-diffusive motion (diffusivity coefficient, α<1) within the experimental timescale (**Fig. 5b, c**). With aging, these condensates exhibit a progressive lowering of the crossover frequency and increase in the storage (G’) and loss (G’’) moduli, suggesting an increase in the network elasticity (**Fig. 5d; Supplementary Fig. 17**). Surprisingly, L-Arg-treated condensates at 2 hours of age were found to be more viscoelastic relative to that of the untreated condition (**Fig. 5b, c**). At 4 hours, these condensates exhibit a substantial increase in viscoelasticity (**Fig. 5d; Supplementary Fig. 17**). These data suggest a surprising role of L-Arg in strengthening intra-condensate networking interactions, otherwise referred to as percolation^87, 88^. Optical tweezer-based active fusion measurements of SynTag-Tau condensate show a consensus with nanorheology measurements (**Fig. 5b vs. Supplementary Fig. 18a, Fig. 1h**) in that L-Arg leads to a slowdown in condensate fusion speed in a dose-dependent manner. In agreement with these results, FRAP measurements at longer timescales revealed progressively slower recovery traces for L-Arg-treated condensates (**Supplementary Fig. 18b**). We also tested the effect of L-Glu treatment on condensate viscoelasticity (**Supplementary Fig. 19**). We observed that it enhances condensate viscoelasticity similarly to L-Arg; however, it is worth noting that this enhancement might stem from faster fibril assembly (**Supplementary Fig. 16**) rather than elevated condensate metastability. Further systematic investigation will be required to clearly distinguish these mechanisms. In total, measurements of condensate material properties reveal that L-Arg suppresses condensate-to-fibril transition via enhancement of condensate viscoelasticity.

**Figure 5.**
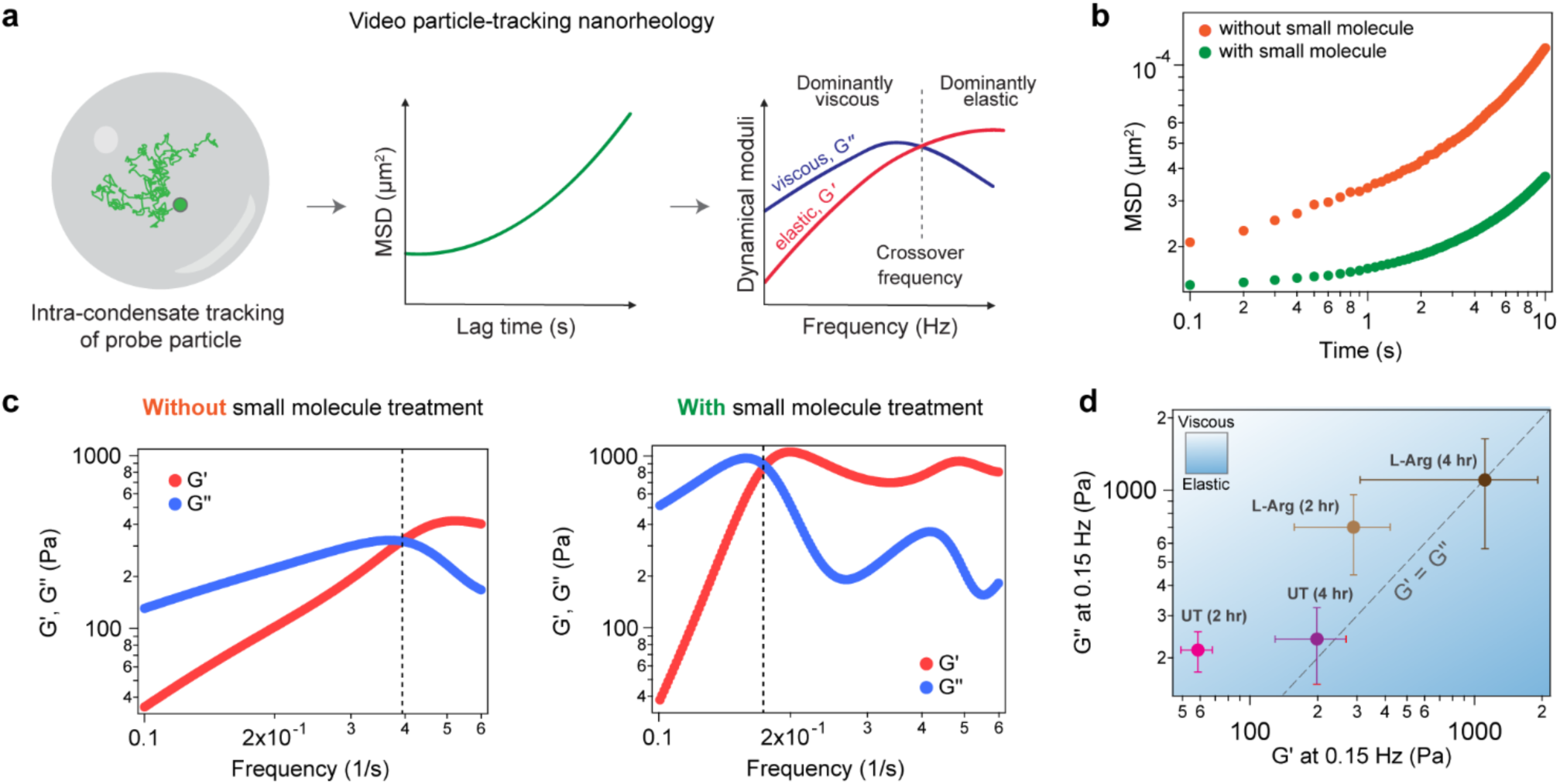
Small molecule-mediated enhancement of condensate viscoelasticity. **(a)** Schematic of VPT nanorheology measurements using 200 nm probe particles passively embedded within Tau condensates. **(b)** Representative mean squared displacement (MSD) measurements of probe particles inside Tau condensates with or without small molecule treatment (2 mM L-Arg). The measurements were conducted 2 hours after sample preparation. **(c)** Representative dynamical moduli of Tau condensates either in the absence (left) or presence (right) of 2 mM L-Arg. The measurements were conducted 2 hours after sample preparation. The dashed line represents the crossover frequency. **(d)** Reports a diagram-of-states for untreated (UT) and small molecule-treated (L-Arg) Tau condensates at the indicated sample age, based on the measured moduli at 0.15 Hz. The experimentally determined data are represented as mean values with standard deviations. The composition of the samples used here is 40 μM Tau in buffer containing 10 mM HEPES (pH 7.4), 50 mM NaCl, 0.1 mM EDTA, and 2 mM DTT along with 7.5% PEG8000, and 6.25 μΜ heparin. Wherever applicable, the L-Arg concentration used in experiments is 2 mM. The sample size for all these measurements is 3-5 condensates from three independent replicates.

### Selective inhibition of condensate-to-fibril transition preserves condensate biochemical activity

The effect of L-Arg showing protective effects against the condensate-to-fibril transition of SynTag-Tau motivated us to test its role in MT assembly (**Fig. 2**). We observe that, similar to untreated SynTag-Tau condensates, the L-Arg-treated condensates can assemble MTs with comparable efficiency up to 5 hours of physical aging (**Fig. 6a, b; Supplementary Fig. 20a**). After the 5-hour mark, these trends start to diverge. The untreated SynTag-Tau condensates progressively lose their MT assembly activity as seen by the reduced MT formation (**Fig. 2d, e)**. In contrast, the L-Arg-treated SynTag-Tau condensates (1 mM and 2 mM L-Arg) show a steady trend in MT assembly activity on the same timescale without appreciable decline. Particularly, at the 16-hour time point, the untreated SynTag-Tau condensates completely lose activity, whereas the L-Arg-treated condensates continue to retain their function (**Fig. 6a, b; Supplementary Fig. 20a**). Tubulin partitioning experiments conducted under similar assay conditions revealed that L-Arg treatment is associated with homogeneous tubulin partitioning into condensates at all time points, which is otherwise perturbed as a function of condensate age (**Supplementary Fig. 12**). Moreover, FRAP measurements reveal marginally slower mobility of tubulin with L-Arg treatment, relative to the untreated condition, likely owed to the small molecule-induced enhancement in condensate viscoelasticity (**Supplementary Fig. 20b; Fig. 5**). Nonetheless, the outcome of tubulin assembly does not appear to be dramatically affected by small molecule-mediated changes in condensate material state (**Fig. 6b**). With our overall observations, we posit that nucleation of amyloid assemblies at the SynTag-Tau condensate interface results in the accumulation of tubulin at the condensate interface, hindering MT assembly. In the presence of L-Arg, fibril nucleation at the condensate interface is inhibited, allowing favorable partitioning of tubulin to the condensate core and facilitating tubulin polymerization (**Fig. 6c**).

**Fig. 6.**
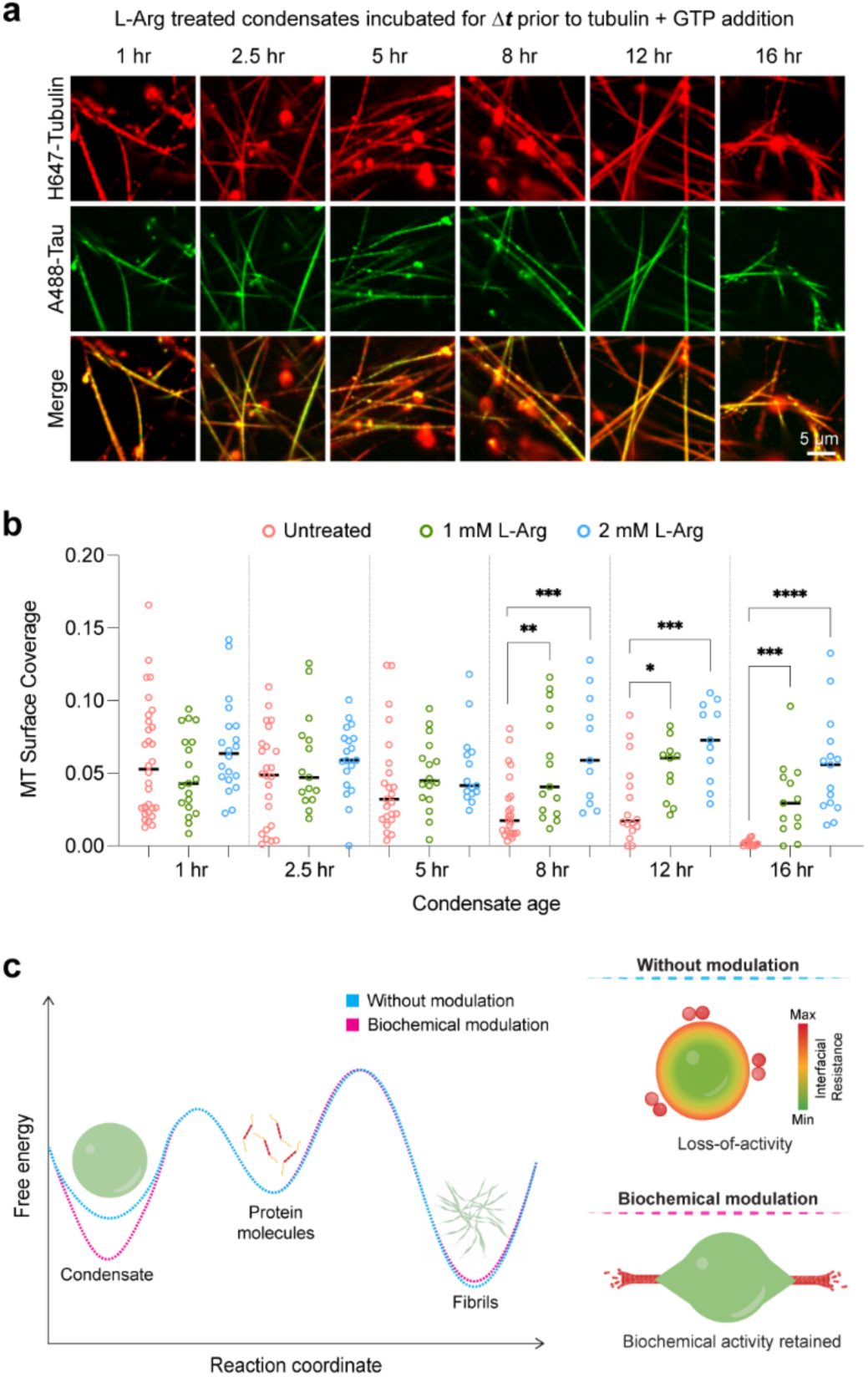
Selective inhibition of condensate age-dependent fibril formation preserves condensate biochemical activity. **(a)** MT polymerization assay of SynTag-Tau condensates treated with 1 mM L-Arg at various time points. **(b)** A comparison of MT surface coverage as a function of condensate age between untreated SynTag-Tau condensates (Fig. 2e) and L-Arg treated SynTag-Tau condensates (images corresponding to 2 mM condition are shown in **Supplementary Fig. 20a**). Horizontal black lines represent the median value. The individual data points represent 11-30 measurements, based on three independent replicates. Statistical significance was determined using an unpaired two-sided Student’s *t*-test between the MT surface coverages of untreated condensates (orange) and either 1 mM L-Arg treated condensates (green) or 2 mM L-Arg treated condensates (blue) (* means p<0.05, ** means p<0.01, *** means p<0.001, **** means p<0.0001). The associated *P* values are shown from left to right: 0.0028, 0.0003, 0.0264, 0.0006, 0.0002, and 0.00009. **(c)** Schematic representing the proposed model for the biochemical modulation of condensate metastability governing protein condensate-to-fibril transition and its impact on the condensate biochemical activity. The composition of samples used in panels (a) and (b) is 12 μΜ SynTag-Tau protein in buffer containing 80 mM PIPES (pH 6.9), 2 mM MgCl_2_, 0.5 mM EGTA, and 2 mM DTT, along with 5% PEG8000 crowder. The tubulin concentration used here is 5 μM, and the concentration of GTP is 1 mM. The concentrations of Atto488-labeled SynTag-Tau and HiLyte647-labeled tubulin are 250 nM and 500 nM, respectively. These experiments were independently repeated three times.

## Conclusion

Previous reports have suggested that liquid-like biomolecular condensates may act as hubs for the aggregation of disease-linked proteins^20, 22, 89, 90^. An important question in this regard is whether the driving forces for phase separation and amyloid fibril formation are the same or merely overlapping. If condensates were to be crucibles for amyloid fibril formation^91^, one would speculate that physicochemical factors that stabilize the condensed phase of proteins would result in a commensurate acceleration in fibril formation. Using sequence engineering of the prion-like domain of hnRNPA1, recent reports have independently shown the separation of functions—the separability of sequence features that drive phase separation and fibril formation^17, 24^. In our studies, we demonstrate an orthogonal approach to the separation of function paradigm by utilizing small molecules to modulate the material properties of protein condensates. Although protein sequence engineering has been a common way to dissect the interplay between condensate formation and aging into stable solids, such approaches are not feasible to apply in disease models. Our results provide a proof-of-principle that the free energy landscape for condensate-to-fibril transition can be selectively modulated by small molecules (**Fig. 6c**), thereby preserving condensate biochemical activity.

What is the mechanism of L-Arg’s action in inhibiting Tau condensate-to-fibril transition? Firstly, we find that the Tau condensate interface plays a critical role in nucleating fibril formation, consistent with other protein systems that show similar aging behavior^17, 79, 80^. Therefore, the mechanism of L-Arg’s action may rest, in part, on its ability to prevent amyloid nucleation at the Tau condensate interface, a process that leads to a crust-like solid shell formation in aged condensates^92^. Inhibition of the solidification of the interface of Tau condensates would enable favorable partitioning of tubulin to the condensate core, allowing MT assembly. Secondly, L-Arg strengthens the metastable viscoelastic network of Tau condensates, thereby delaying the onset of the condensate-to-fibril transition. A similar mechanism has recently been proposed in the instance of Tau condensates treated with methylene blue^57^, a long-studied drug candidate in clinical trials of Tauopathies^93^. Within the intracellular environment, L-Arg concentration is in the lower millimolar range^85, 86^, similar to that used in our study, which leads us to speculate that L-Arg may act as a natural regulator of aberrant protein phase transitions, similar to the proposed role of ATP as a biological hydrotrope^26^. In contrast, we found that some other small molecule metabolites, such as L-Asp and L-Glu, enhance the Tau liquid-to-solid transition. These findings inspire a hypothesis that the cellular metabolic state itself is a critical regulator of protein phase separation and aggregation in living systems. Under diverse conditions, cells can produce a variety of metabolites that can dynamically tune the aging dynamics of biomolecular condensates.

The current study employs a unique strategy of introducing a synthetic prionogenic sequence tag to full-length Tau protein, which lowers the nucleation barrier of the condensate-to-fibril transition without perturbing condensate formation and its biochemical activity. Our findings are in corroboration with growing lines of evidence that have directly correlated the impact of disease mutations to the weakening of condensate metastability and, inversely, global stabilization of the fibrillar solid phase^12, 24, 94^. These observations warrant further investigations on the molecular grammars of condensate metastability and aging timescales^95^. The designed Tau protein system used in the study provided valuable insight into the effect of material properties and physical aging on condensate biochemical activity.

In summary, protein condensates are metastable fluids that must overcome a free-energy barrier to transition to fibrils, which is a more thermodynamically stable state (**Fig. 6c**). Our study reinforces the framework of separation of functions of protein phase separation and physical aging into irreversible solids. We experimentally demonstrated that these two processes can be decoupled by small molecules that selectively enhance condensate metastability and thereby inhibit condensate-to-amyloid transition. Our results may inspire strategies to selectively target protein pathological fibrillization without perturbing their functional phase separation.

## Data availability

All data are available in the manuscript or the supplementary materials.

## Code availability

The codes for nanorheology-related analyses are available on GitHub (see github.com/BanerjeeLab-repertoire/Decoupling-Phase-Separation-and-Fibrillization-Preserves-Condensate-Biochemical-Activity).

## Acknowledgments

This work was supported by the US National Institutes of Health (NIH) through grants R03 AG070510 (to P.R.B.), R35 GM138186 (to P.R.B.), and the St. Jude Children’s Research Collaborative on the Biology and Biophysics of RNP Granules (to P.R.B.). We are grateful for financial support from the Welch Foundation F-2008-20220331, the Chan Zuckerberg Initiative #2021-236087, and the National Science Foundation (NSF) #2146549 to S.H.P., and C.M.J. is grateful for support from NSF (DGE-1610403) and NIH T32 EB007507. We would like to gratefully acknowledge Dr. Ibraheem Alshareedah, currently at Los Alamos National Laboratory, as well as Paul Dewan, currently at Harvard University, for technical support during the development phase of the study. Also, the authors greatly appreciate the critical feedback from the group members of the Banerjee laboratory, as well as Drs. R. Pappu and T. Mittag. Any opinions, findings, conclusions, or recommendations expressed in this material are those of the author(s) and do not necessarily reflect the views of the NSF, NIH, or any other bodies.

## Author contributions

Conceptualization: P.R.B. and T.S.M.; Methodology: P.R.B., T.S.M., A.S., S.S., S.H.P., C.M.J., and C.N.; Investigation: P.R.B., T.S.M., A.S., S.S., S.H.P., C.M.J., and B.H.G.; Resources: P.R.B. and S.H.P.; Writing – original draft and revisions: P.R.B. and T.S.M.; Writing – reviewing and editing: P.R.B., T.S.M., A.S., S.S., S.H.P., C.M.J.; Funding acquisition: P.R.B. and S.H.P.

## References

1. Martin EW, Holehouse AS, Peran I, Farag M, Incicco JJ, Bremer A, et al. Valence and patterning of aromatic residues determine the phase behavior of prion-like domains. Science 2020, 367(6478): 694–699.

2. Ginell GM, Holehouse AS. An Introduction to the Stickers-and-Spacers Framework as Applied to Biomolecular Condensates. In: Zhou H-X, Spille J-H, Banerjee PR (eds). Phase-Separated Biomolecular Condensates: Methods and Protocols. Springer US: New York, NY, 2023, pp 95–116.

3. Pappu RV, Cohen SR, Dar F, Farag M, Kar M. Phase Transitions of Associative Biomacromolecules. Chemical Reviews 2023, 123(14): 8945–8987.

4. Dignon GL, Best RB, Mittal J. Biomolecular Phase Separation: From Molecular Driving Forces to Macroscopic Properties. Annual Review of Physical Chemistry 2020, 71(Volume 71, 2020): 53-75.

5. Rekhi S, Garcia CG, Barai M, Rizuan A, Schuster BS, Kiick KL, et al. Expanding the molecular language of protein liquid–liquid phase separation. Nature Chemistry 2024, 16(7): 1113–1124.

6. Brangwynne CP, Eckmann CR, Courson DS, Rybarska A, Hoege C, Gharakhani J, et al. Germline P Granules Are Liquid Droplets That Localize by Controlled Dissolution/Condensation. Science 2009, 324(5935): 1729–1732.

7. Alshareedah I, Moosa MM, Pham M, Potoyan DA, Banerjee PR. Programmable viscoelasticity in protein-RNA condensates with disordered sticker-spacer polypeptides. Nat Commun 2021, 12(6620): 1–14.

8. Jawerth L, Fischer-Friedrich E, Saha S, Wang J, Franzmann T, Zhang X, et al. Protein condensates as aging Maxwell fluids. Science 2020, 370(6522): 1317-1323.

9. Alshareedah I, Borcherds WM, Cohen SR, Singh A, Posey AE, Farag M, et al. Sequence-specific interactions determine viscoelasticity and ageing dynamics of protein condensates. Nature Physics 2024.

10. Kato M, Han TW, Xie S, Shi K, Du X, Wu LC, et al. Cell-free formation of RNA granules: low complexity sequence domains form dynamic fibers within hydrogels. Cell 2012, 149(4): 753–767.

11. Molliex A, Temirov J, Lee J, Coughlin M, Kanagaraj AP, Kim HJ, et al. Phase Separation by Low Complexity Domains Promotes Stress Granule Assembly and Drives Pathological Fibrillization. Cell 2015, 163(1): 123–133.

12. Patel A, Lee HO, Jawerth L, Maharana S, Jahnel M, Hein MY, et al. A Liquid-to-Solid Phase Transition of the ALS Protein FUS Accelerated by Disease Mutation. Cell 2015, 162(5): 1066–1077.

13. St George-Hyslop P, Lin JQ, Miyashita A, Phillips EC, Qamar S, Randle SJ, et al. The physiological and pathological biophysics of phase separation and gelation of RNA binding proteins in amyotrophic lateral sclerosis and fronto-temporal lobar degeneration. Brain Res 2018, 1693(Pt A): 11–23.

14. Lin Y, Protter DS, Rosen MK, Parker R. Formation and Maturation of Phase-Separated Liquid Droplets by RNA-Binding Proteins. Mol Cell 2015, 60(2): 208–219.

15. Zbinden A, Pérez-Berlanga M, De Rossi P, Polymenidou M. Phase Separation and Neurodegenerative Diseases: A Disturbance in the Force. Dev Cell 2020, 55(1): 45–68.

16. Murakami T, Qamar S, Lin JQ, Schierle GS, Rees E, Miyashita A, et al. ALS/FTD Mutation-Induced Phase Transition of FUS Liquid Droplets and Reversible Hydrogels into Irreversible Hydrogels Impairs RNP Granule Function. Neuron 2015, 88(4): 678–690.

17. Linsenmeier M, Faltova L, Morelli C, Capasso Palmiero U, Seiffert C, Küffner AM, et al. The interface of condensates of the hnRNPA1 low-complexity domain promotes formation of amyloid fibrils. Nat Chem 2023, 15(10): 1340–1349.

18. Zhou X, Sumrow L, Tashiro K, Sutherland L, Liu D, Qin T, et al. Mutations linked to neurological disease enhance self-association of low-complexity protein sequences. Science 2022, 377(6601): eabn5582.

19. Alberti S, Dormann D. Liquid–Liquid Phase Separation in Disease. Annual Review of Genetics 2019, 53(Volume 53, 2019): 171-194.

20. Alberti S, Carra S. Quality Control of Membraneless Organelles. Journal of Molecular Biology 2018, 430(23): 4711–4729.

21. Alberti S, Gladfelter A, Mittag T. Considerations and Challenges in Studying Liquid-Liquid Phase Separation and Biomolecular Condensates. Cell 2019, 176(3): 419–434.

22. Alberti S, Hyman AA. Biomolecular condensates at the nexus of cellular stress, protein aggregation disease and ageing. Nat Rev Mol Cell Biol 2021, 22(3): 196–213.

23. Babinchak WM, Surewicz WK. Liquid–Liquid Phase Separation and Its Mechanistic Role in Pathological Protein Aggregation. J Mol Biol 2020, 432(7): 1910–1925.

24. Das T, Zaidi FK, Farag M, Ruff KM, Mahendran TS, Singh A, et al. Tunable metastability of condensates reconciles their dual roles in amyloid fibril formation. Molecular Cell 2025, 85(11): 2230–2245.e2237.

25. Zhang P, Fan B, Yang P, Temirov J, Messing J, Kim HJ, et al. Chronic optogenetic induction of stress granules is cytotoxic and reveals the evolution of ALS-FTD pathology. eLife 2019, 8: e39578.

26. Patel A, Malinovska L, Saha S, Wang J, Alberti S, Krishnan Y, et al. ATP as a biological hydrotrope. Science 2017, 356(6339): 753-756.

27. Choi K-J, Tsoi PS, Moosa MM, Paulucci-Holthauzen A, Liao S-CJ, Ferreon JC, et al. A Chemical Chaperone Decouples TDP-43 Disordered Domain Phase Separation from Fibrillation. Biochemistry 2018, 57(50): 6822–6826.

28. Qamar S, Wang G, Randle SJ, Ruggeri FS, Varela JA, Lin JQ, et al. FUS Phase Separation Is Modulated by a Molecular Chaperone and Methylation of Arginine Cation-π Interactions. Cell 2018, 173(3): 720–734.e715.

29. Gu J, Liu Z, Zhang S, Li Y, Xia W, Wang C, et al. Hsp40 proteins phase separate to chaperone the assembly and maintenance of membraneless organelles. Proceedings of the National Academy of Sciences 2020, 117(49): 31123–31133.

30. Linsenmeier M, Hondele M, Grigolato F, Secchi E, Weis K, Arosio P. Dynamic arrest and aging of biomolecular condensates are modulated by low-complexity domains, RNA and biochemical activity. Nat Commun 2022, 13(1): 3030.

31. Hofweber M, Hutten S, Bourgeois B, Spreitzer E, Niedner-Boblenz A, Schifferer M, et al. Phase Separation of FUS Is Suppressed by Its Nuclear Import Receptor and Arginine Methylation. Cell 2018, 173(3): 706–719.e713.

32. Babinchak WM, Dumm BK, Venus S, Boyko S, Putnam AA, Jankowsky E, et al. Small molecules as potent biphasic modulators of protein liquid-liquid phase separation. Nature Communications 2020, 11(1): 5574.

33. Wheeler RJ, Lee HO, Poser I, Pal A, Doeleman T, Kishigami S, et al. Small molecules for modulating protein driven liquid-liquid phase separation in treating neurodegenerative disease. bioRxiv 2019: 721001.

34. Freibaum BD, Messing J, Nakamura H, Yurtsever U, Wu J, Kim HJ, et al. Identification of small molecule inhibitors of G3BP-driven stress granule formation. Journal of Cell Biology 2024, 223(3).

35. Klein IA, Boija A, Afeyan LK, Hawken SW, Fan M, Dall’Agnese A, et al. Partitioning of cancer therapeutics in nuclear condensates. Science 2020, 368(6497): 1386-1392.

36. Mitrea DM, Mittasch M, Gomes BF, Klein IA, Murcko MA. Modulating biomolecular condensates: a novel approach to drug discovery. Nature Reviews Drug Discovery 2022, 21(11): 841–862.

37. Patel A, Mitrea D, Namasivayam V, Murcko MA, Wagner M, Klein IA. Principles and functions of condensate modifying drugs. Front Mol Biosci 2022, 9: 1007744.

38. Dada ST, Toprakcioglu Z, Cali MP, Röntgen A, Hardenberg MC, Morris OM, et al. Pharmacological inhibition of α-synuclein aggregation within liquid condensates. Nature Communications 2024, 15(1): 3835.

39. Ambadi Thody S, Clements HD, Baniasadi H, Lyon AS, Sigman MS, Rosen MK. Small-molecule properties define partitioning into biomolecular condensates. Nature Chemistry 2024, 16(11): 1794–1802.

40. Ray S, Singh N, Kumar R, Patel K, Pandey S, Datta D, et al. α-Synuclein aggregation nucleates through liquid-liquid phase separation. Nat Chem 2020, 12(8): 705–716.

41. Lee VM, Goedert M, Trojanowski JQ. Neurodegenerative tauopathies. Annu Rev Neurosci 2001, 24: 1121–1159.

42. Zhang Y, Wu K-M, Yang L, Dong Q, Yu J-T. Tauopathies: new perspectives and challenges. Molecular Neurodegeneration 2022, 17(1): 28.

43. Goedert M, Eisenberg DS, Crowther RA. Propagation of Tau Aggregates and Neurodegeneration. Annual review of neuroscience 2017, 40: 189–210.

44. Wang Y, Mandelkow E. Tau in physiology and pathology. Nat Rev Neurosci 2016, 17(1): 22–35.

45. Götz J, Halliday G, Nisbet RM. Molecular Pathogenesis of the Tauopathies. Annu Rev Pathol: Mech Dis 2019, 14(1): 239–261.

46. Hernández-Vega A, Braun M, Scharrel L, Jahnel M, Wegmann S, Hyman BT, et al. Local Nucleation of Microtubule Bundles through Tubulin Concentration into a Condensed Tau Phase. Cell Rep 2017, 20(10): 2304–2312.

47. Tan R, Lam AJ, Tan T, Han J, Nowakowski DW, Vershinin M, et al. Microtubules gate tau condensation to spatially regulate microtubule functions. Nature Cell Biology 2019, 21(9): 1078–1085.

48. Ash PEA, Lei S, Shattuck J, Boudeau S, Carlomagno Y, Medalla M, et al. TIA1 potentiates tau phase separation and promotes generation of toxic oligomeric tau. Proceedings of the National Academy of Sciences 2021, 118(9): e2014188118.

49. Kanaan NM, Hamel C, Grabinski T, Combs B. Liquid-liquid phase separation induces pathogenic tau conformations in vitro. Nat Commun 2020, 11.

50. Wegmann S, Eftekharzadeh B, Tepper K, Zoltowska KM, Bennett RE, Dujardin S, et al. Tau protein liquid–liquid phase separation can initiate tau aggregation. The EMBO journal 2018, 37(7).

51. Boyko S, Surewicz K, Surewicz WK. Regulatory mechanisms of tau protein fibrillation under the conditions of liquid–liquid phase separation. Proc Natl Acad Sci USA 2020, 117(50): 31882–31890.

52. Hochmair J, Exner C, Franck M, Dominguez-Baquero A, Diez L, Brognaro H, et al. Molecular crowding and RNA synergize to promote phase separation, microtubule interaction, and seeding of Tau condensates. EMBO J 2022: e108882.

53. Meisl G, Knowles TPJ, Klenerman D. Mechanistic Models of Protein Aggregation Across Length-Scales and Time-Scales: From the Test Tube to Neurodegenerative Disease. Front Neurosci 2022, 16: 909861.

54. Meisl G, Hidari E, Allinson K, Rittman T, DeVos SL, Sanchez JS, et al. In vivo rate-determining steps of tau seed accumulation in Alzheimer’s disease. Sci Adv 2021, 7(44): eabh1448.

55. Mandelkow E-M, Mandelkow E. Biochemistry and cell biology of tau protein in neurofibrillary degeneration. Cold Spring Harbor perspectives in medicine 2012, 2(7): a006247.

56. Lin Y, Fichou Y, Zeng Z, Hu NY, Han S. Electrostatically Driven Complex Coacervation and Amyloid Aggregation of Tau Are Independent Processes with Overlapping Conditions. ACS Chem Neurosci 2020, 11(4): 615–627.

57. Huang Y, Wen J, Ramirez L-M, Gümüşdil E, Pokhrel P, Man VH, et al. Methylene blue accelerates liquid-to-gel transition of tau condensates impacting tau function and pathology. Nature Communications 2023, 14(1): 5444.

58. Savastano A, Flores D, Kadavath H, Biernat J, Mandelkow E, Zweckstetter M. Disease-Associated Tau Phosphorylation Hinders Tubulin Assembly within Tau Condensates. Angew Chem Int Ed 2021, 60(2): 726–730.

59. Chuang H-Y, He R-Y, Huang Y-A, Hsu W-T, Cheng Y-J, Guo Z-R, et al. Engineered droplet-forming peptide as photocontrollable phase modulator for fused in sarcoma protein. Nature Communications 2024, 15(1): 5686.

60. Piroska L, Fenyi A, Thomas S, Plamont M-A, Redeker V, Melki R, et al. α-Synuclein liquid condensates fuel fibrillar α-synuclein growth. Science Advances 2023, 9(33): eadg5663.

61. Khan T, Kandola TS, Wu J, Venkatesan S, Ketter E, Lange JJ, et al. Quantifying Nucleation In Vivo Reveals the Physical Basis of Prion-like Phase Behavior. Molecular Cell 2018, 71(1): 155–168.e157.

62. Goehler H, Dröge A, Lurz R, Schnoegl S, Chernoff YO, Wanker EE. Pathogenic Polyglutamine Tracts Are Potent Inducers of Spontaneous Sup35 and Rnq1 Amyloidogenesis. PLoS One 2010, 5(3): e9642.

63. Chandramowlishwaran P, Sun M, Casey KL, Romanyuk AV, Grizel AV, Sopova JV, et al. Mammalian amyloidogenic proteins promote prion nucleation in yeast. J Biol Chem 2018, 293(9): 3436–3450.

64. Toombs JA, Petri M, Paul KR, Kan GY, Ben-Hur A, Ross ED. De novo design of synthetic prion domains. Proc Natl Acad Sci U S A 2012, 109(17): 6519–6524.

65. Michiels E, Liu S, Gallardo R, Louros N, Mathelié-Guinlet M, Dufrêne Y, et al. Entropic Bristles Tune the Seeding Efficiency of Prion-Nucleating Fragments. Cell Rep 2020, 30(8): 2834–2845.e2833.

66. Alberti S, Halfmann R, King O, Kapila A, Lindquist S. A systematic survey identifies prions and illuminates sequence features of prionogenic proteins. Cell 2009, 137(1): 146–158.

67. Halfmann R, Alberti S, Krishnan R, Lyle N, O’Donnell CW, King OD, et al. Opposing effects of glutamine and asparagine govern prion formation by intrinsically disordered proteins. Mol Cell 2011, 43(1): 72–84.

68. Abramson J, Adler J, Dunger J, Evans R, Green T, Pritzel A, et al. Accurate structure prediction of biomolecular interactions with AlphaFold 3. Nature 2024, 630(8016): 493-500.

69. Mahendran TS, Bremer A, Singh A, Basalla J, Marzahn MR, Mittag T, et al. BPS2025 - Network strength and interface concentration in multicomponent condensates determine their abilities to suppress fibril formation. Biophysical Journal 2025, 124(3): 197a.

70. Goldschmidt L, Teng PK, Riek R, Eisenberg D. Identifying the amylome, proteins capable of forming amyloid-like fibrils. Proceedings of the National Academy of Sciences 2010, 107(8): 3487–3492.

71. Jennings CM, Markel AC, Domingo MJE, Miller KS, Bayer CL, Parekh SH. Collagen organization and structure in FBLN5-/- mice using label-free microscopy: implications for pelvic organ prolapse. Biomed Opt Express 2024, 15(5): 2863–2875.

72. Kwan AC, Duff K, Gouras GK, Webb WW. Optical visualization of Alzheimer’s pathology via multiphoton-excited intrinsic fluorescence and second harmonic generation. Opt Express 2009, 17(5): 3679–3689.

73. Chatterjee S, Kan Y, Brzezinski M, Koynov K, Regy RM, Murthy AC, et al. Reversible Kinetic Trapping of FUS Biomolecular Condensates. Adv Sci (Weinh) 2022, 9(4): e2104247.

74. Najafi S, Lin Y, Longhini AP, Zhang X, Delaney KT, Kosik KS, et al. Liquid–liquid phase separation of Tau by self and complex coacervation. Protein Sci 2021, 30(7): 1393–1407.

75. Ukmar-Godec T, Hutten S, Grieshop MP, Rezaei-Ghaleh N, Cima-Omori M-S, Biernat J, et al. Lysine/RNA-interactions drive and regulate biomolecular condensation - Nature Communications. Nat Commun 2019, 10(2909): 1–15.

76. Kaur T, Alshareedah I, Wang W, Ngo J, Moosa MM, Banerjee PR. Molecular Crowding Tunes Material States of Ribonucleoprotein Condensates. Biomolecules 2019, 9(2): 71.

77. Wei M-T, Elbaum-Garfinkle S, Holehouse AS, Chen CC-H, Feric M, Arnold CB, et al. Phase behaviour of disordered proteins underlying low density and high permeability of liquid organelles. Nature Chemistry 2017, 9(11): 1118–1125.

78. Takarai H, Yasuda T, Sakumichi N, Sakai T. Partitioning Law of Polymer Chains into Flexible Polymer Networks. arXiv preprint arXiv:250505254 2025.

79. Emmanouilidis L, Bartalucci E, Kan Y, Ijavi M, Pérez ME, Afanasyev P, et al. A solid beta-sheet structure is formed at the surface of FUS droplets during aging. Nature Chemical Biology 2024.

80. Shen Y, Chen A, Wang W, Shen Y, Ruggeri FS, Aime S, et al. The liquid-to-solid transition of FUS is promoted by the condensate surface. Proc Natl Acad Sci U S A 2023, 120(33): e2301366120.

81. Farag M, Cohen SR, Borcherds WM, Bremer A, Mittag T, Pappu RV. Condensates formed by prion-like low-complexity domains have small-world network structures and interfaces defined by expanded conformations. Nat Commun 2022, 13(1): 7722.

82. Ruff KM, Dar F, Pappu RV. Polyphasic linkage and the impact of ligand binding on the regulation of biomolecular condensates. Biophys Rev (Melville) 2021, 2(2): 021302.

83. Mathieu C, Pappu RV, Taylor JP. Beyond aggregation: Pathological phase transitions in neurodegenerative disease. Science 2020, 370(6512): 56-60.

84. Mahendran TS, Wadsworth GM, Singh A, Gupta R, Banerjee PR. Homotypic RNA clustering accompanies a liquid-to-solid transition inside the core of multi-component biomolecular condensates. Nature Chemistry 2025, 17(8): 1236–1246.

85. Joshi MS, Ferguson TB, Johnson FK, Johnson RA, Parthasarathy S, Lancaster JR. Receptor-mediated activation of nitric oxide synthesis by arginine in endothelial cells. Proceedings of the National Academy of Sciences 2007, 104(24): 9982–9987.

86. Baydoun AR, Emery PW, Pearson JD, Mann GE. Substrate-dependent regulation of intracellular amino acid concentrations in cultured bovine aortic endothelial cells. Biochem Biophys Res Commun 1990, 173(3): 940–948.

87. Mittag T, Pappu RV. A conceptual framework for understanding phase separation and addressing open questions and challenges. Molecular Cell 2022, 82(12): 2201–2214.

88. Harmon TS, Holehouse AS, Rosen MK, Pappu RV. Intrinsically disordered linkers determine the interplay between phase separation and gelation in multivalent proteins. eLife 2017, 6: e30294.

89. Boeynaems S, Alberti S, Fawzi NL, Mittag T, Polymenidou M, Rousseau F, et al. Protein Phase Separation: A New Phase in Cell Biology. Trends Cell Biol 2018, 28(6): 420–435.

90. Lipiński WP, Visser BS, Robu I, Fakhree MAA, Lindhoud S, Claessens MMAE, et al. Biomolecular condensates can both accelerate and suppress aggregation of α-synuclein. Science Advances 2022, 8(48): eabq6495.

91. Li YR, King OD, Shorter J, Gitler AD. Stress granules as crucibles of ALS pathogenesis. Journal of Cell Biology 2013, 201(3): 361–372.

92. Biswas S, Potoyan DA. Molecular Drivers of Aging in Biomolecular Condensates: Desolvation, Rigidification, and Sticker Lifetimes. PRX Life 2024, 2(2): 023011.

93. Wang L, Bharti, Kumar R, Pavlov PF, Winblad B. Small molecule therapeutics for tauopathy in Alzheimer’s disease: Walking on the path of most resistance. European Journal of Medicinal Chemistry 2021, 209: 112915.

94. Garcia-Cabau C, Bartomeu A, Tesei G, Cheung KC, Pose-Utrilla J, Picó S, et al. Mis-splicing of a neuronal microexon promotes CPEB4 aggregation in ASD. Nature 2025, 637(8045): 496-503.

95. Wake N, Weng SL, Zheng T, Wang SH, Kirilenko V, Mittal J, et al. Expanding the molecular grammar of polar residues and arginine in FUS prion-like domain phase separation and aggregation. bioRxiv 2024.

