## Supplementary Information for "Decoupling Phase Separation and Fibrillization Preserves Activity of Biomolecular Condensates"

### Materials and Methods

#### Protein purification

Codon-optimized constructs of wild-type Tau (2N4R isoform) and designed variants used in this study were synthesized and cloned into a pET His6 N10 TEV LIC vector by GenScript. SynTag-Tau protein was expressed and purified under denaturing conditions from the E. coli BL21 strain (New England Biolabs). Initially, E. coli cells were grown to $OD_{600}$ of 0.6-0.8 and subsequently induced with 0.1 mM IPTG at 37 °C for 2 hours. After pelleting, the cells were resuspended in a lysis buffer composed of 50 mM phosphate buffer (pH 6.4), 1 M KCl, 10 mM imidazole, 2 mM DTT, 6 M urea, and a protease inhibitor cocktail (Thermo Fisher Scientific). The resuspended lysate solution was subject to ultrasonication followed by ultracentrifugation at 37,500xg for 30 mins at 30 °C. The extracted supernatant from the previous step was introduced to a packed Ni-NTA column at 4 °C (5 ml volume; Thermo Fisher Scientific) prewashed with the lysis buffer. After several washing steps, the proteins were eluted from the column using an elution buffer composed of 50 mM phosphate buffer (pH 6.4), 2 M KCl, 100 mM imidazole, 2 mM DTT, and 3 M urea. The elutes were pooled and introduced to a pre-equilibrated Zeba$^{TM}$ spin desalting column (Thermo Fisher Scientific) following the manufacturer's instructions to get rid of residual imidazole and for storage in a buffer composed of 50 mM phosphate buffer (pH 6.4), 2 M NaCl, 2 mM DTT, and 3 M urea. This step was repeated twice to ensure efficient desalting. The resulting protein stock was tested for nucleic acid contamination, and the protein concentration was estimated by NanoDrop 1C$^{TM}$ spectrophotometer.

In the case of wild-type (WT) Tau (2N4R isoform) with 6xHis tag, the protein was expressed and purified under native conditions from the E. coli BL21 strain (New England Biolabs). Initially, E. coli cells were grown to $OD_{600}$ of 0.6-0.8 and subsequently induced with 0.1 mM IPTG at 37 °C for 2.5 hours. After pelleting, the cells were resuspended in a lysis buffer composed of 50 mM MES (pH 6.0), 300 mM NaCl, 10 mM imidazole, 2 mM DTT, 1 mM EDTA, 1 mM phenylmethylsulfonyl fluoride (PMSF, Sigma Aldrich), and a protease inhibitor cocktail (Thermo Fisher Scientific). The resuspended lysate solution was subject to ultrasonication followed by ultracentrifugation at 37,500xg for 30 mins at 4 °C. The extracted supernatant was then incubated with Ni-NTA resin (pre-equilibrated with the lysis buffer) overnight at 4 °C (5 ml volume; Thermo Fisher Scientific). The protein was eluted from the Ni-column after several wash steps using an elution buffer composed of 50 mM MES (pH 6.0), 300 mM NaCl, 250 mM imidazole, 2 mM DTT, 1 mM EDTA, and 0.3 mM PMSF. Protein fractions containing 6xHis-tagged Tau protein were incubated with TurboTEV protease (Eton Bioscience) overnight at 4°C. The proteolysis reaction volume was loaded onto another Ni-column to remove the free 6xHis tag, non-cleaved protein, and TEV protease. The supernatant containing unbound Tau protein was diluted in a buffer containing 50 mM MES, pH 6.0, and 2 mM DTT and then loaded onto a Capto HiRes S 5/50 cation exchange column (Cytiva) for ion-exchange chromatography. Tau protein was eluted from the column by a step gradient of 0 to 1 M NaCl. Purified protein in storage buffer containing 50 mM MES (pH 6.0), 500 mM NaCl, and 2 mM DTT was tested for nucleic acid contamination, and the protein concentration was estimated by NanoDrop 1C$^{TM}$ spectrophotometer.

Finally, SDS-PAGE analysis was conducted to test the quality of all the purified proteins prior to flash-freezing the protein aliquots and storage at -80 °C. Site-specific fluorescence labeling of proteins was accomplished following the manufacturer's instructions. For the sequence details of WT and Tau variants used in this study, refer to **Supplementary Table 1**.

#### Preparation of protein condensate samples

After thawing an aliquot of purified Tau protein sample at room temperature, desalting using an equilibrated Zeba$^{TM}$ spin desalting column (Thermo Fisher Scientific) was conducted, following the manufacturer's instructions, using a buffer composed of 10 mM HEPES (pH 7.5), 200 mM NaCl, 0.1 mM EDTA, and 2 mM DTT. The choice of this buffer composition for a majority of the experiments conducted in the study is based on the observation that it provides conditions for favorable protein phase separation into large condensate sizes that are amenable to experimental interrogation, and for clear demarcation of condensate interface from their interior (Supplementary Fig. 7). The previous step was repeated twice to ensure efficient desalting. The resulting protein was concentrated, and its concentration was measured using a NanoDrop 1C spectrophotometer. The final sample for experiments was then prepared by diluting the concentrated protein solution in the same buffer without NaCl. Next, PEG8000 (Sigma Aldrich) was added at a desired concentration (specified in the figure legends) to trigger phase separation. Whenever applicable, small molecules were introduced into the sample buffer during the protein dilution step. Small molecules, including L-arginine, D-arginine, L-arginine ethyl ester, L-lysine, L-proline, L-aspartic acid, L-glutamic acid, adenosine triphosphate, α-dansyl-L-arginine, dansyl chloride, and trimethylamine N-oxide, were procured from Sigma Aldrich; urea and guanidinium hydrochloride were procured from Thermo Fisher Scientific. Much of the small molecule stocks were prepared with water, except for α-dansyl chloride and α-dansyl-L-arginine, which were dissolved in water spiked with DMSO; the final concentration of DMSO in condensate samples is less than 1%. Multivalent polymers such as penta-lysine, deca-lysine, and deca-arginine were procured from GenScript, whereas spermine and spermidine were procured from Sigma Aldrich. The stocks of these multivalent polymers were prepared and introduced into the sample buffer, similar to the small molecules. In Thioflavin T (ThT; Thermo Fisher Scientific) based experiments, ThT at a final concentration of 50 μM was introduced during the buffer dilution step. Pipetting was carried out subsequently to ensure sufficient mixing of all components in the sample to form protein condensates. The final sample was sandwiched between a tween 20-coated coverslip and a glass slide separated by three layers of double-sided tape. The remaining void was filled with mineral oil to seal the sample chamber to prevent evaporation. Fluorescence imaging was performed either using a Q2 laser scanning confocal microscope (ISS Inc., 63× objective) or a Lumicks (63× objective) C-trap confocal microscope, or a Zeiss LSM710 (63× objective) confocal microscope. Andor Dragonfly 600 confocal platform (Oxford Instruments), equipped with an Andor iXon Ultra EMCCD camera and a Leica HC PL APO 63x/1.40-0.60 oil objective, was used for generating z-stack fluorescence images.

The preparation of wild-type Tau condensates was carried out similarly to that of SynTag-Tau and related variants, with the exception of including 6.25 mM heparin (Santa Cruz Biotechnology) in the sample buffer (unless specified otherwise) prior to induction of phase separation by the addition of PEG8000. The sample was then placed inside an incubator set at 37 °C. These conditions were shown previously to enhance the physical aging of Tau condensates[1].

### Fluorescence recovery after photobleaching (FRAP)

Protein condensate samples prepared in a Tween 20-coated sample chamber were kept on a confocal microscope stage (Lumicks C-trap) and imaged with a 60× water immersion objective of numerical aperture 1.3. Imaging steps of either Atto488-labeled SynTag-Tau or HiLyte647-labeled tubulin were accomplished using the respective excitation laser line. Selection of the region of interest (ROI) for reference, bleaching, and continuous imaging for monitoring fluorescence recovery was done using the Bluelake software (Lumicks, https://lumicks.github.io/bluelake-api/2.4.0/index.html). The bleaching ROI dimensions were kept consistent for all condensate samples. Each condensate sample was bleached for at least 5 frames at maximum laser power to achieve optimal bleach depth, with a dwell time of 150 ms per pixel. Recovery of fluorescence was recorded for a maximum of 300 seconds after the bleaching steps. The images from the FRAP experiments were analyzed using Fiji[2] (version 1.54f) to quantify the fluorescence recovery as a function of time. The intensities were normalized and corrected as described previously[3]. Succinctly, the normalization accounts for a correction for background intensity after the bleaching step as well as the photo-fading that follows, using the following expression, wherein $I_{correct}(t)$ is the intensity of the bleached ROI at time $t$ corrected for photofading and $I_{correct}(0)$ is the value of intensity before the start of the bleach step (reference ROI used for quantifying the bleaching depth):

$$I_{normalized}(t) = \frac{I_{correct}(t) - \min(I_{correct})}{I_{correct}(0) - \min(I_{correct})} \quad (1)$$

In total, 4 to 5 fluorescence recovery traces representative of three independent measurements were gathered and averaged.

### Second-harmonic generation (SHG) imaging

SHG measurements were made using a Nd:YAG microchip laser generating nanosecond pulses with a repetition rate of approximately 1 MHz at 1064 nm (Leukos Opera HP, Leukos). A 100×, 0.85 NA objective (LCPLN100XIR, Olympus) focused the beam to the sample plane, and the transmission SHG signal was collected with a 20×, 0.4 NA objective (M-20X, MKS Newport). The signal was measured with a spectrometer (IsoPlane 160, Teledyne Princeton Instruments) and a back-illuminated, deep-depletion CCD (Blaze 1340 x 400 HS, Teledyne Princeton Instruments). The CCD integration time per pixel was 1000 ms. Samples were stage-scanned with a pixel spacing of 0.3 μm per pixel.

### Optical tweezer-mediated coalescence assay

Optical trap-induced fusion of protein condensates was accomplished using a Lumicks C-trap optical tweezer system equipped with a dual trap and a laser scanning confocal fluorescence microscope, with a 60x water immersion objective of numerical aperture 1.3, following a protocol described previously[4, 5]. In brief, for a typical condensate fusion experiment, two condensates were trapped using a 1064 nm laser with the minimum power setting (~ 100 μW) to minimize laser-induced heating effects. Typically, the 1064 nm laser used in the assay provides a power of 1 mW at the objective (measured at the exit of the laser light from the objective). For the experiments, a power of 10% of 1 mW, which corresponds to ~100 μW, was used. This power is sufficient to trap droplets using the dual-trap optical tweezer system and perform droplet fusion assays[4]. The trapping of condensates relies on the refractive index mismatch between the condensate dense phase and the surrounding dilute phase. After trapping, Trap 2 was kept fixed, and Trap 1 was moved towards Trap 2 at a constant velocity of 100 nm/s. The trap is set to maintain this velocity until the fusion process concludes, allowing the final droplet to relax to a spherical or equilibrium shape. A bright-field camera was used to take time-lapse images of the droplet fusion event at 15 frames per second.

### Broadband coherent anti-Stokes Raman (BCARS) hyperspectral imaging

For structural analysis of SynTag-Tau conformers within condensates or fibrils, we used an in-house-built, broadband coherent anti-Stokes Raman scattering microscope. For excitation, a Nd:YAG microchip laser generates nanosecond pulses with a repetition rate of approximately 1 MHz at 1064 nm and a broadband supercontinuum that ranges from 1100 – 2400 nm (Leukos Opera HP, Leukos). The beams were focused on the sample plane via a 100×, 0.85 NA objective (LCPLN100XIR, Olympus). In a transmission configuration, the signal was collected with a 20×, 0.4 NA objective (M-20X, MKS Newport). The signal was measured with a spectrometer (IsoPlane 160, Teledyne Princeton Instruments) and a back-illuminated, deep-depletion CCD (Blaze 1340 x 400 HS, Teledyne Princeton Instruments). Curve-fitting measurements were performed by scanning the sample with a pixel spacing of 0.4 μm. The integration time per pixel was 80-150 ms to maximize the signal without saturating the CCD.

### Broadband coherent anti-Stokes Raman (BCARS) data processing

As reported in previous studies, raw BCARS spectra were transformed into Raman-like spectra for quantitative analysis using a modified Kramers-Kronig transform for phase retrieval[6, 7]. Then, a second-order Savisky-Golay with a 151-point (approximately Δ492 $cm^{-1}$) smoothing window was applied to the phase-retrieved spectra to produce the Raman-like spectra.

An average spectrum for phase-separated condensates was extracted using a 3x3 pixel mean (condensate center). Fibril average spectra were calculated as the mean spectra within a drawn region of interest around the fibril. The average Raman-like spectra were subsequently normalized by the max spectral intensity[8] arising from the CH3 vibration located at 2933.4 $cm^{-1}$. BCARS post-processing is performed in Igor Pro 8.4 (WaveMetrics). For consistency, center wavelength dependencies of the spectrometer calibration required a spectral shift of less than or equal to 10 $cm^{-1}$ before amide I curve-fitting. A Gaussian peak was fit to the CH3 peak to identify the center of the CH3 peak. Seven spectral points were linearly interpolated between each measured wavenumber, and the entire spectra shifted by the difference between the CH3 center and 2933.4 $cm^{-1}$.

Amide I decomposition was performed with Igor Pro MultiPeak Fitting 2 functionality, which employs an iterative Levenberg-Marquardt algorithm. The parameters of the Lorentzian peaks for curve-fitting are found in **Supplementary Table 2**. Briefly, we used six peaks[9, 10, 11] within the amide I band (1550 – 1725 $cm^{-1}$): 1644 $cm^{-1}$ for α-helices, 1660 $cm^{-1}$ for random coils, 1673 $cm^{-1}$ for β-sheets, a minor peak for β-turns at 1691 $cm^{-1}$, and two minor peaks for tyrosine ring modes at 1600 $cm^{-1}$ and 1612 $cm^{-1}$.

### Tubulin partitioning and microtubule assembly functional assay

Tubulin and HiLyte647-labeled tubulin were purchased from Cytoskeleton. Partitioning experiments, using HiLyte647-labeled tubulin, were conducted on condensate samples composed of SynTag-Tau protein at 24 μM protein concentration with a PEG 8000 concentration of 10% with the following buffer composition: 10 mM HEPES (pH 7.4), 50 mM NaCl, 0.1 mM EDTA, and 2 mM DTT. The HiLyte647-labeled tubulin was introduced to SynTag-Tau condensates that were allowed to incubate within a microfuge tube for a defined period (as indicated in the text). The concentration of HiLyte647-labeled tubulin is specified in the text. In addition, the partitioning experiments were also performed under sample conditions corresponding to the microtubule polymerization assay (described below). The microtubule polymerization assay was carried out based on methods described previously[12, 13]. For a schematic of the assay, refer to Fig. 2a. Briefly, SynTag-Tau protein solution was buffer-exchanged to general tubulin buffer (Cytoskeleton) consisting of 80 mM PIPES (pH 6.9), 2 mM $MgCl_2$, and 0.5 mM EGTA, supplemented with 2 mM DTT. This buffer composition was specifically used for this assay, as it was shown to support robust and reproducible tubulin polymerization under near-physiological conditions[14]. The addition of PEG8000 to the buffer-exchanged sample at a final concentration of 5% induced the formation of protein condensates. These condensates are allowed to incubate within a microfuge tube for a defined period, as mentioned in the text, before the addition of tubulin and GTP at final concentrations of 5 μM and 1 mM, respectively. 500 nM HiLyte647-labeled tubulin was doped along with unlabeled tubulin for MT polymerization assays. Finally, the sample was prepared for fluorescence microscopy as described in the '*Preparation of protein condensate samples'* method section, with imaging conducted after the sample incubation period of 30 minutes.

### Image analysis to estimate microtubule surface coverage

The analysis of microtubule surface coverage was performed using a custom-made Python-based image analysis program. Initially, raw images were loaded into the analysis program and subjected to median filtering to suppress noise, followed by Richardson-Lucy deconvolution to enhance the resolution and clarity. To further accentuate these structures, gray-scale dilation and erosion were applied alongside contrast stretching and Gabor filtering, which specifically highlighted the microtubular objects in the image. Following enhancement, the brightest objects, predominantly condensates, were isolated through an intensity threshold. These objects were then removed using binary morphological operations combined with region labeling, where the eccentricity of each region was calculated to differentiate spherical condensates from microtubule structures. After excluding the primary condensates, the processed images were subjected to another round of thresholding to selectively identify microtubules. This selection was refined by assessing the eccentricity of dimmer blobs, which were likely overlooked in the initial removal step. Such blobs were excluded to avoid falsely conflating condensates with the microtubule structures. Final mask refinement involved additional binary morphological operations to fine-tune the segmentation. The resulting microtubule mask was then measured quantitatively. The final mask was measured, and data, including an overlay image and a data frame of measurements, such as microtubular coverage, were exported for further analysis.

### Frequency-domain fluorescence lifetime imaging (FD-FLIM)

FD-FLIM measurements of protein condensate samples were collected using a Q2 laser scanning confocal microscope (ISS Inc.) equipped with FastFLIM™ modules[15]. For calibration prior to FLIM measurements, Rhodamine110 dye diluted in water, with a fluorescence lifetime of 4.0 ns, was used. Fitting of the phase delay and modulation ratio of the acquired FD-FLIM data of a given condensate sample at digital modulation frequencies ranging from 20 to 100 MHz was used to estimate the fluorescence lifetime distribution of Atto488-labeled SynTag-Tau. Subsequently, this was used to map the distribution of fluorescence lifetimes in the condensate images. Generation of the corresponding fluorescence lifetime histograms of condensate samples at different imaging time points, as well as thresholding, was carried out using the ISS VistaVision software. Gaussian fitting of the fluorescence lifetime histograms was accomplished using GraphPad Prism 10.

### Image analysis for estimating partition coefficient

Condensate samples used for partition coefficient analysis were imaged either using a Lumicks C-trap confocal microscope (60×) or a Zeiss LSM710 confocal microscope (63×). The concentration of labeled proteins/molecules, as well as the settings used for the respective excitation laser lines, was kept consistent across samples. Fluorescence images collected for partition coefficient image analysis were analyzed using Fiji[2] to segment individual condensates and estimate the mean intensity of each condensate (dense phase), referred to as $I_{dense}$. To estimate the background mean fluorescence intensity (dilute phase) referred to as $I_{dilute}$, three regions away from the condensates were selected at random. The partition coefficient (k) is finally estimated by dividing the dense phase mean intensity and the dilute phase mean intensity, k= $I_{dense}$/ $I_{dilute}$.

### Estimation of Thioflavin T (ThT) fluorescence intensity

Condensate samples were prepared following a protocol as outlined in the '*Preparation of protein condensate samples*' method section. The samples were imaged either using a Lumicks C-Trap confocal microscope or an ISS Q2 laser scanning confocal microscope, wherein, for each set of experiments, the microscope used and the acquisition settings were kept consistent. The acquired fluorescence images were quantified using Fiji to estimate the mean ThT fluorescence intensity, with normalization to the background ThT fluorescence intensity. These ThT measurements were performed either in the absence or presence of a small molecule to assess whether the small molecule treatment leads to enhanced or reduced fibril formation kinetics using ThT fluorescence intensity as a proxy.

### Video particle tracking (VPT) based nanorheology

VPT-based nanorheology measurements were conducted on wild-type Tau condensates following the protocol outlined in a previous report[16]. In brief, we utilized 200 nanometer-sized carboxylate-modified polystyrene beads (yellow-green fluorescent tracers; FluoSpheres™, Invitrogen) introduced into the sample buffer (0.0005 % solids) prior to induction of Tau phase separation, along with heparin to trigger Tau condensate aging. The remainder of the sample preparation was carried out as prescribed in the '*Preparation of protein condensate samples*' method section. The protein concentration used for these measurements is higher relative to the typical concentration used for other experiments, as reported in the main text, to allow for the formation of larger condensate volumes, which is favorable for tracking a higher number of fluorescent tracers for nanorheology measurements. To determine the mean-squared displacements of the tracers, the Trackmate[17] plugin of Fiji was used.

The acquired trajectories were subjected to correction for potential drift by subtracting from the center of mass (COM) trajectory, which was estimated using particle velocity as shown below:

$$\boldsymbol{X_{COM}}(k) = \boldsymbol{X_0} + \sum_{j=0}^{k} \frac{1}{N_j} \sum_{i=1}^{N} \boldsymbol{v_{i,j}} \tag{2}$$

Here, $k$ represents the frame number at which $\boldsymbol{X_{COM}}$ is calculated, $j$ represents the individual frame, and $N_j$ represents the number of particles of frame $j$. $\boldsymbol{X_0}$ represents the center of the mass vector of the first frame. Subsequently, after the correction of the trajectories for drifting, the mean squared displacement (MSD) is estimated as follows:

$$MSD(\tau) = \langle \boldsymbol{R}(t+\tau) - \boldsymbol{R}(t) \rangle_{t,N} \tag{3}$$

Here, $\tau$ represents the lag time and $\boldsymbol{R}$ represents the position vector of the particle. Utilizing various lag times $\tau$, the MSD was estimated based on the trajectories of the particles, with the use of in-house made

Python scripts, which are publicly available on GitHub (github.com/BanerjeeLab-repertoire/Decoupling-Phase-Separation-and-Fibrillization-Preserves-Condensate-Biochemical-Activity). Next, the ensemble averaged MSD was estimated using measurements from approximately 20–100 individual probe particles. Fitting of the MSD[4, 18] was accomplished using:

$$MSD(\tau) = 4D\tau^{\alpha} + N \quad (4)$$

Here, $D$ represents the diffusion coefficient of the particles and $\alpha$ represents the diffusivity exponent, which signifies the nature of the particle diffusion (normal or sub-diffusive) inside the condensates.

With each sample, about 20 to 200 trajectories were acquired from a range of 3 to 5 condensates across three independently prepared samples. In order to deduce the condensate aging dynamics in the presence or absence of small molecules (L-Arg), VPT nanorheology was conducted at various time points during physical aging.

### Estimation of dynamical moduli from nanorheology measurements

The viscoelastic properties of Tau condensates were estimated from the MSD measurements, utilizing in-house developed Python scripts (github.com/BanerjeeLab-repertoire/Decoupling-Phase-Separation-and-Fibrillization-Preserves-Condensate-Biochemical-Activity). The protocol followed to estimate the dynamical moduli from MSDs is described in a previous report[19]. In brief, the compliance $j(t)$ was calculated from the MSDs $< \Delta r^2(\tau) >$ estimated from VPT nanorheology measurements described in the previous section using the following equation:

$$j(t) = \frac{3\pi a}{n_d k_B T} < \Delta r^2(\tau) > \quad (5)$$

Here, $a$ represents the probe particle radius, $n_d$ = 2 represents the dimension of the probe particle trajectories based on which the MSDs were estimated, $k_B$ represents the Boltzmann constant, and $T$ represents the absolute temperature at which VPT nanorheology experiments were done.

The complex shear modulus can be estimated using a Fourier transform, which relates the complex shear modulus and the compliance via a convolution integral, as shown below:

$$\int_0^{\tau} G(t) j(\tau - t) dt = \tau \quad (6)$$

$$G^*(\omega) = \frac{1}{i\omega \hat{j}(\omega)}. \quad (7)$$

Following the methodology proposed by Evans, R. et al.[20], the complex shear modulus was estimated. Using discrete experimental data points $(t_i, j_i)$ of the estimated compliance from Eq. (5), the frequency-space shear modulus $G^*(\omega)$ is estimated following the equation below:

$$\frac{i\omega}{G^*(\omega)} = i\omega J(0) + \frac{\left(1 - e^{-i\omega t_1}\right)\left(J_1 - J(0)\right)}{t_1} + \frac{e^{-i\omega t_N}}{\eta} + \sum_{k=2}^{N} \left(\frac{J_k - J_{k-1}}{t_k - t_{k-1}}\right)\left(e^{-i\omega t_{k-1}} - e^{-i\omega t_k}\right) \quad (8)$$

The MSD data that were measured using nanorheology were oversampled using a cubic spline for estimation. The remainder of the parameters required in Eq. (8) for estimating the shear modulus can be acquired as follows. $j(0)$ is acquired through extrapolation of the experimental data for compliance $t = 0$ using a linear fit of the initial four data points and $\eta$ using a linear fit of the final ten data points following the relation, $\eta = 1/j(t)$.

### Estimation of dense phase area fraction

Confocal fluorescence images visualized with Atto488-labeled SynTag-Tau were used as input for dense phase area fraction estimation. The imaging was conducted after all condensates had settled on the coverslip. Random frames were imaged to ensure an accurate representation of the surface area. For quantification, thresholding was performed using Fiji in order to isolate condensates from the background (dilute phase), yielding a binarized image displaying only the highlighted areas occupied by condensates. Using the 'Analyze Particles' function of Fiji, individual condensate sizes ($\mu m^2$) were determined. Subsequently, the fraction of cumulative area occupied by condensates with respect to the total area of image acquisition, set by image acquisition parameters, was estimated. The dense phase area fraction estimations were conducted at different conditions, including varying buffer ionic strength or small molecule concentrations, to assess their impact on phase separation.

### Coverslip treatment for microscopy

The experiments of this study used Tween20-coated coverslips. The coverslip cleaning procedure prior to surface coating involves treatment with 2% Hellmanex solution for 2 hours, followed by rinsing with MilliQ water six times. Subsequently, coverslips were treated with 20% Tween 20 solution (v/v) for 30 minutes. Further, these coverslips were rinsed with MilliQ six times and dried with compressed air. Lastly, the coverslips were dried in an oven set to 40 °C with overnight incubation. The coverslips were later stored at room temperature for use in experiments.

### Software

Fiji[2] (version 1.54f) was used for processing and analyzing microscopy images. A custom Python script was used for plotting VPT nanorheology data and estimation of dynamical moduli. The scripts are available on GitHub (github.com/BanerjeeLab-repertoire/Decoupling-Phase-Separation-and-Fibrillization-Preserves-Condensate-Biochemical-Activity). Trackmate (v7.14) was used for nanorheology-related analysis. For all other plots, GraphPad Prism 10 (v10.4.1) was used. Marvin Sketch (Chemaxon) was used to visualize chemical structures of small molecules. Adobe Illustrator (2024) was used for preparing figures and schematics. Igor Pro 8.4 was used for BCARS data processing. Bluelake (v1.6.11) was used for collecting fluorescence images as well as for performing FRAP and optical tweezer-induced condensate fusion measurements with the Lumicks C-Trap microscope. ZEN (SP5 2012 Black) was used for acquiring fluorescence images with the LSM 710 laser scanning confocal microscope. Vistavision (v4.2) was used for acquiring fluorescence images as well as FLIM measurements with the ISS Q2 laser scanning confocal microscope. Fusion Dragonfly software (v2.6) was used for acquiring fluorescence images with the Andor Dragonfly 600 confocal platform. UCSF ChimeraX[21] (v1.7) was used for visualizing AlphaFold3[22] predicted structures of Tau protein variants. ZipperDB was used for estimating fibrillization propensity scores[23]. DrugBank was used for generating the predicted pKa values of small molecules[24]. PLAAC[25] was used to generate prion scores of protein variants.

### Supplementary Figures

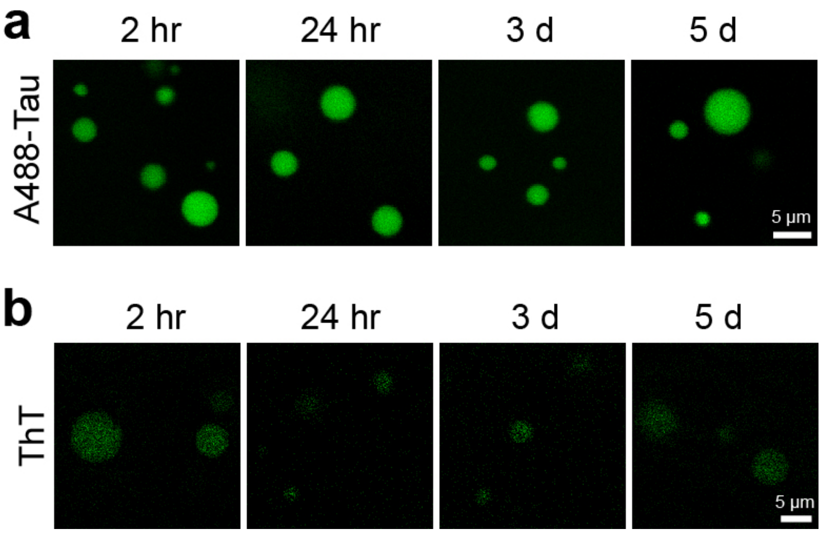


**Supplementary Figure 1**. (a) Wild-type Tau forms phase-separated condensates that do not age to form mesoscale fibrils under quiescent cofactor-free conditions. (b) Tau condensates show an absence of Thioflavin T (ThT) staining as a function of time, indicative of a lack of emergent fibrillar structures under similar conditions as (a). The composition of the samples is 25 µM Tau in buffer containing 10 mM HEPES (pH 7.5), 50 mM NaCl, 0.1 mM EDTA, and 2 mM DTT with 7.5% PEG 8000. Wherever applicable, the concentration of Atto488-labeled Tau is 250 nM, and the concentration of ThT is 50 µM. Each experiment was independently repeated three times.

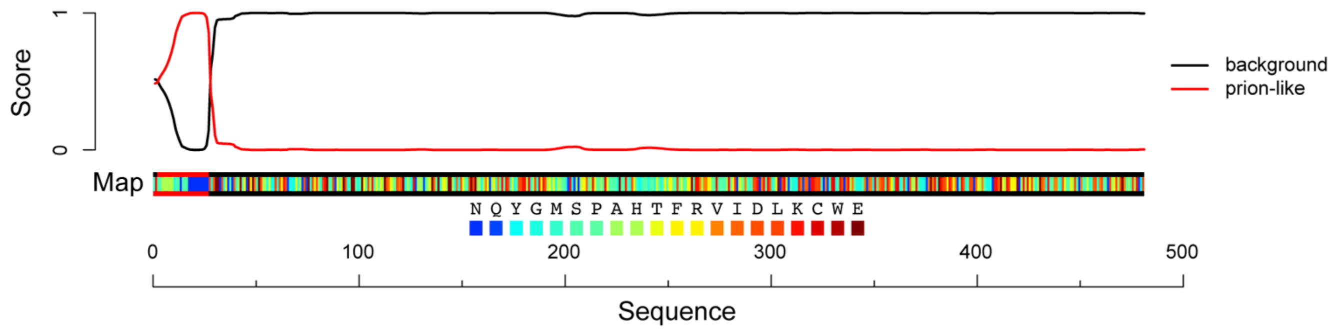


```
1   MKSSHHHHHHGSSNSSSNNNNNNNNNNNLGIEENLYFQSNIMAEPRQEFEVMEDHAGTYGLGDRKDQGGYTMHQDQEGDTDAGLKESPLQTPTEDGSEEPG
101 SETSDAKSTPTAEDVTAPLVDEGAPGKQAAAQPHTEIPEGTTAEEAGIGDTPSLEDEAAGHVTQARMVSKSKDGTGSDDKKAKGADGKTKIATPRGAAPP
201 GQKGQANATRIPAKTPPAPKTPPSSGEPPKSGDRSGYSSPGSPGTPGSRSRTPSLPTPPTREPKKVAVVRTPPKSPSSAKSRLQTAPVPMPDLKNVKSKI
301 GSTENLKHQPGGGKVQIINKKLDLSNVQSKCGSKDNIKHVPGGGSVQIVYKPVDLSKVTSKCGSLGNIHHKPGGGQVEVKSEKLDFKDRVQSKIGSLDNI
401 THVPGGGNKKIETHKLTFRENAKAKTDHGAEIVYKSPVVSGDTSPRHLSNVSSTGSIDMVDSPQLATLADEVSASLAKQGL
```

**Supplementary Figure 2.** PLAAC[25] score of SynTag-Tau shows that the amino-terminal SynTag (sequence: MKSSHHHHHHGSSNSSSNNNNNNNNNNNLGIEENLYFQSNI) is prionogenic (highlighted in red), with the highest weighting contributed by the N-rich sequence block, which is colored in dark blue within the sequence map.

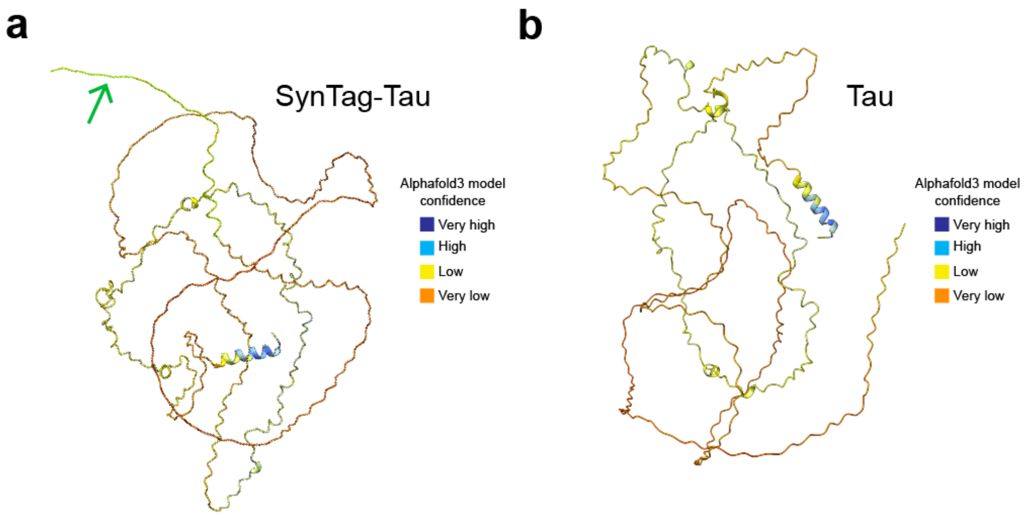


**Supplementary Figure 3**. Alphafold3[22] predicted structure of (a) SynTag-Tau (also shown in Fig. 1c) and (b) wild-type (untagged) Tau protein visualized using ChimeraX[21]. The green arrow in (a) points to the green-color highlighted protein segment corresponding to the SynTag comprised of 40 amino acid residues.

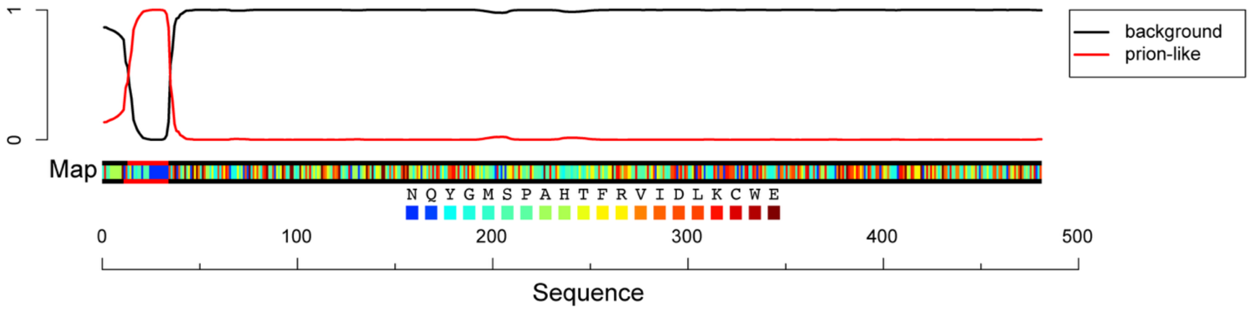


1 MKSSHHHHHHENLYFQSGSSNSSSNNNNNNNNNNLGIENIMAEPRQEFEVMEDHAGTYGLGDRKDQGGYTMHQDQEGDTDAGLKESPLQTPTEDGSEEPG
101 SETSDAKSTPTAEDVTAPLVDEGAPGKQAAAQPHTEIPEGTTAEEAGIGDTPSLEDEAAGHVTQARMVSKSKDGTGSDDKKAKGADGKTKIATPRGAAPP
201 GQKGQANATRIPAKTPPAPKTPPSSGEPPKSGDRSGYSSPGSPGTPGSRSRTPSLPTPPTREPKKVAVVRTPPKSPSSAKSRLQTAPVPMPDLKNVKSKI
301 GSTENLKHQPGGGKVQIINKKLDLSNVQSKCGSKDNIKHVPGGGSVQIVYKPVDLSKVTSKCGSLGNIHHKPGGGQVEVKSEKLDFKDRVQSKIGSLDNI
401 THVPGGGNKKIETHKLTFRENAKAKTDHGAEIVYKSPVVSGDTSPRHLSNVSSTGSIDMVDSPQLATLADEVSASLAKQGL

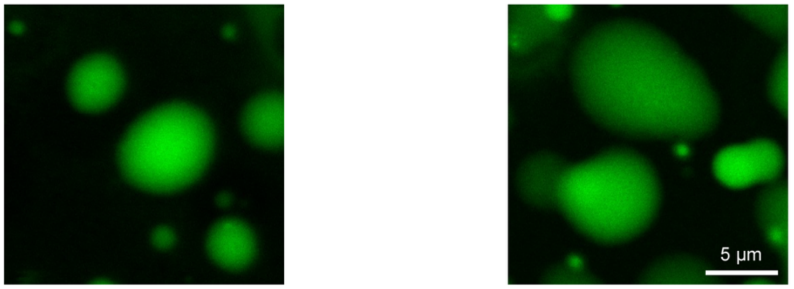


2 hr 5 d

**Supplementary Figure 4**. (top) Sequence perturbations that reduce the PLAAC score[25] of the prionogenic tag in SynTag-Tau without changing its overall amino acid composition prevent the physical aging of condensates into mesoscale fibrillar solids (bottom). Also, see **Supplementary Table 1** for the complete sequence details of the shuffled SynTag-Tau variant used here. The composition of the sample is 12 μM shuffled SynTag-Tau in buffer containing 10 mM HEPES (pH 7.4), 50 mM NaCl, 0.1 mM EDTA, and 2 mM DTT along with 7.5% PEG 8000 crowder. The concentration of Atto488-labeled shuffled SynTag-Tau is 250 nM. This experiment was independently repeated three times.

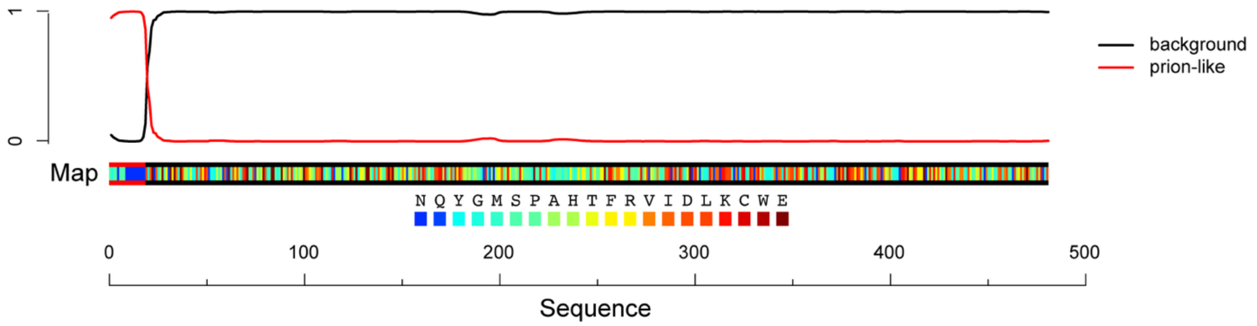


1 SGSSNSSSNNNNNNNNNNNLGIENIMAEPRQEFEVMEDHAGTYGLGDRKDQGGYTMHQDQEGDTDAGLKESPLQTPTEDGSEEPGSETSDAKSTPTAEDVT
101 APLVDEGAPGKQAAAQPHTEIPEGTTAEEAGIGDTPSLEDEAAGHVTQARMVSKSKDGTGSDDKKAKGADGKTKIATPRGAAPPGQKGQANATRIPAKTP
201 PAPKTPPSSGEPPKSGDRSGYSSPGSPGTPGSRSRTPSLPTPPTREPKKVAVVRTPPKSPSSAKSRLQTAPVPMPDLKNVKSKIGSTENLKHQPGGGKVQ
301 IINKKLDLSNVQSKCGSKDNIKHVPGGGSVQIVYKPVDLSKVTSKCGSLGNIHHKPGGGQVEVKSEKLDFKDRVQSKIGSLDNITHVPGGGNKKIETHKL
401 TFRENAKAKTDHGAEIVYKSPVVSGDTSPRHLSNVSSTGSIDMVDSPQLATLADEVSASLAKQGL

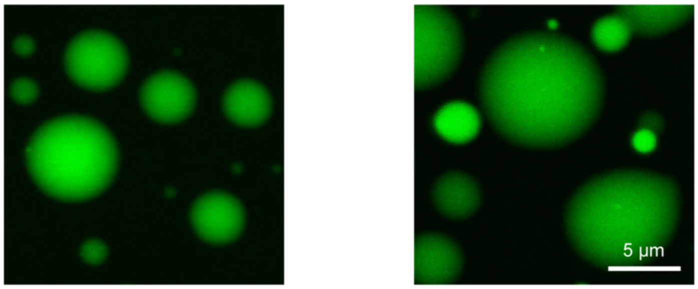


2 hr 5 d

**Supplementary Figure 5**. Truncating the prionogenic tag sequence in SynTag-Tau to the Asn(N)-rich segment, which corresponds to the major weighting in the PLAAC score (top), is insufficient to accelerate the condensate-to-fibril transition (below). Refer to **Supplementary Table 1** for sequence details of the truncated SynTag-Tau variant used here. The composition of the sample is 12 μM truncated SynTag-Tau in buffer containing 10 mM HEPES (pH 7.4), 50 mM NaCl, 0.1 mM EDTA, and 2 mM DTT, along with 7.5% PEG 8000 crowder. The concentration of Atto488-labeled truncated SynTag-475 Tau is 250 nM. This experiment was independently repeated three times.

a

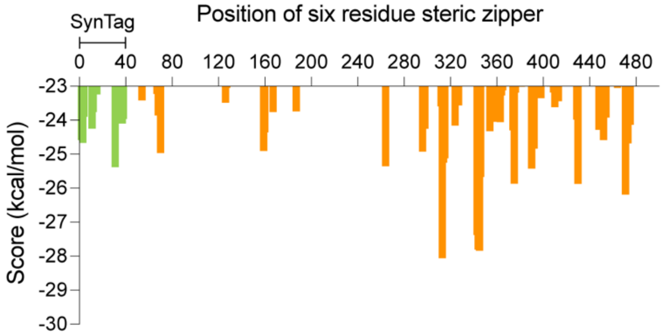


b

| Rank | Steric zipper position | Score (kcal/mol) |
|---|---|---|
| 1 | 313 | -28.05909 |
| 2 | 345 | -27.8337 |
| 3 | 344 | -27.8078 |
| 4 | 343 | -27.37531 |
| 5 | 471 | -26.18819 |
| 6 | 430 | -25.87429 |
| 7 | 375 | -25.86617 |
| 8 | 346 | -25.66898 |
| 9 | 390 | -25.42051 |
| 10 | 31 | -25.38349 |

**Supplementary Figure 6.** (a) Predicted scores of six-residue steric zippers across the SynTag-Tau sequence generated using zipperDB[23], with the Rosetta energy threshold set at -23 kcal/mol. The putative steric zippers in SynTag and the remainder of the full-length Tau sequence are colored green and orange, respectively. (b) Rank order of zipperDB predicted putative steric zippers in SynTag-Tau, corresponding to the analysis done in (a). The row shaded in green indicates a steric zipper confined to the N-terminally attached SynTag, whereas those shaded in orange are confined to the native wild-type Tau sequence.

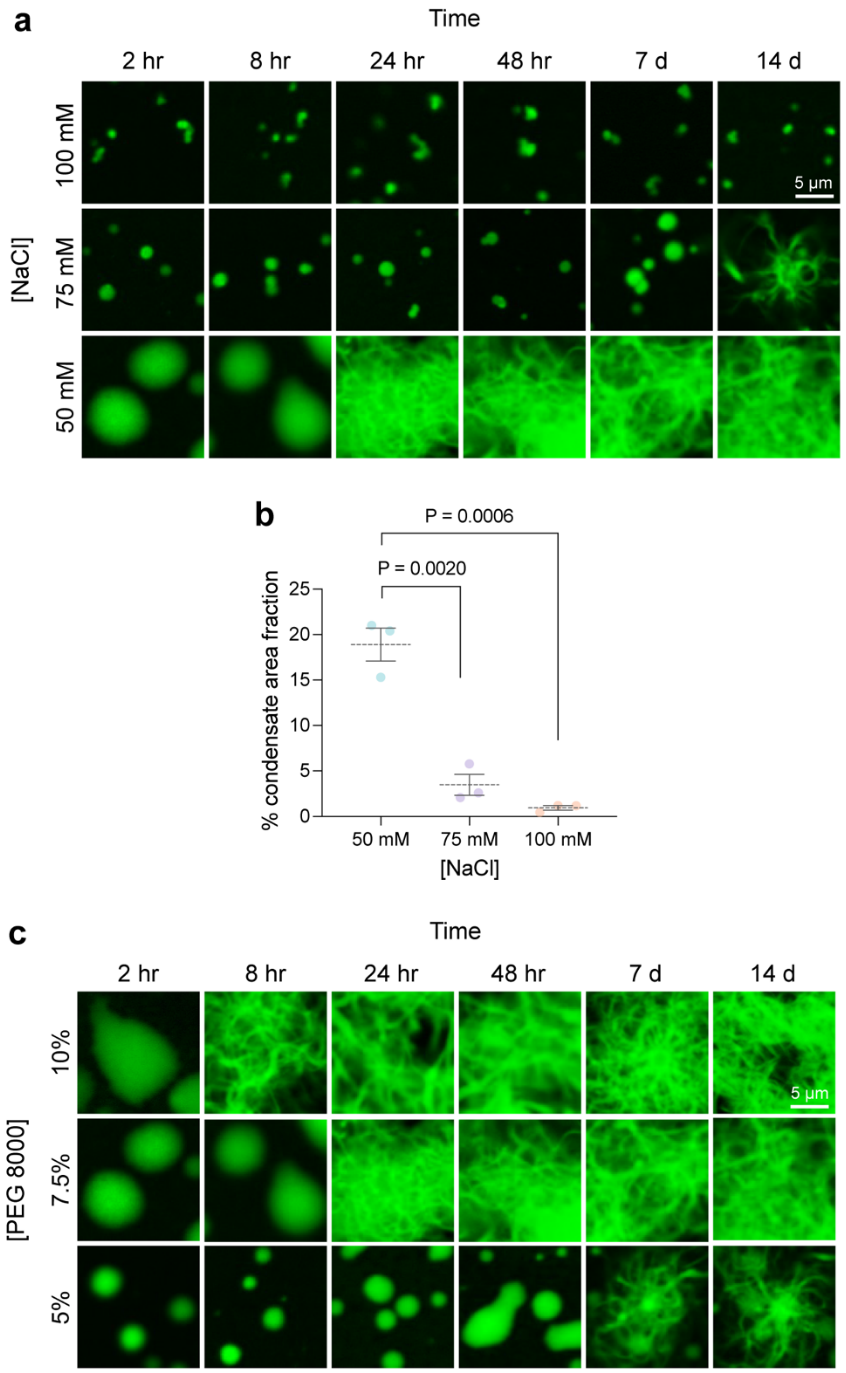


**Supplementary Figure 7**. (a) Effect of modulation of ionic strength through NaCl titration in sample buffer on SynTag-Tau condensate formation and transition to fibrils (d: days). The concentration of SynTag-Tau used here is 12 μM with a fixed PEG8000 crowder concentration of 7.5%, and the concentration of NaCl is specified in the figure. (b) Estimated SynTag-Tau condensate area fraction (expressed in percentages with respect to the total image acquisition area) at different ionic strengths, measured 2 hours after sample preparation. The composition of samples used for these analyses is similar to that described in (a). The dashed line represents the mean, and the error bars represent the standard error of the mean (S.E.M.) based on three independent replicates. Individual points represent data from three independent experiments. Statistical significance was determined using the unpaired two-sided Student's *t*-test. The respective P values are indicated in the plot. (c) Effect of modulation of molecular crowding through PEG8000 titration in sample buffer on SynTag-Tau condensate formation and transition to fibrils.

The concentration of SynTag-Tau used here is 12 μM with a fixed NaCl concentration of 50 mM, and the concentration of PEG8000 crowder is specified in the figure. Please note that the representative '7.5% PEG 8000' data reported here is the same as that shown in the representative '50 mM NaCl' data in panel (a). The buffer composition in all samples is 10 mM HEPES (pH 7.4), 0.1 mM EDTA, and 2 mM DTT, containing NaCl and PEG8000 as indicated. Wherever applicable, the concentration of Atto488-labeled SynTag-Tau is 250 nM. Each experiment was independently repeated at least three times.

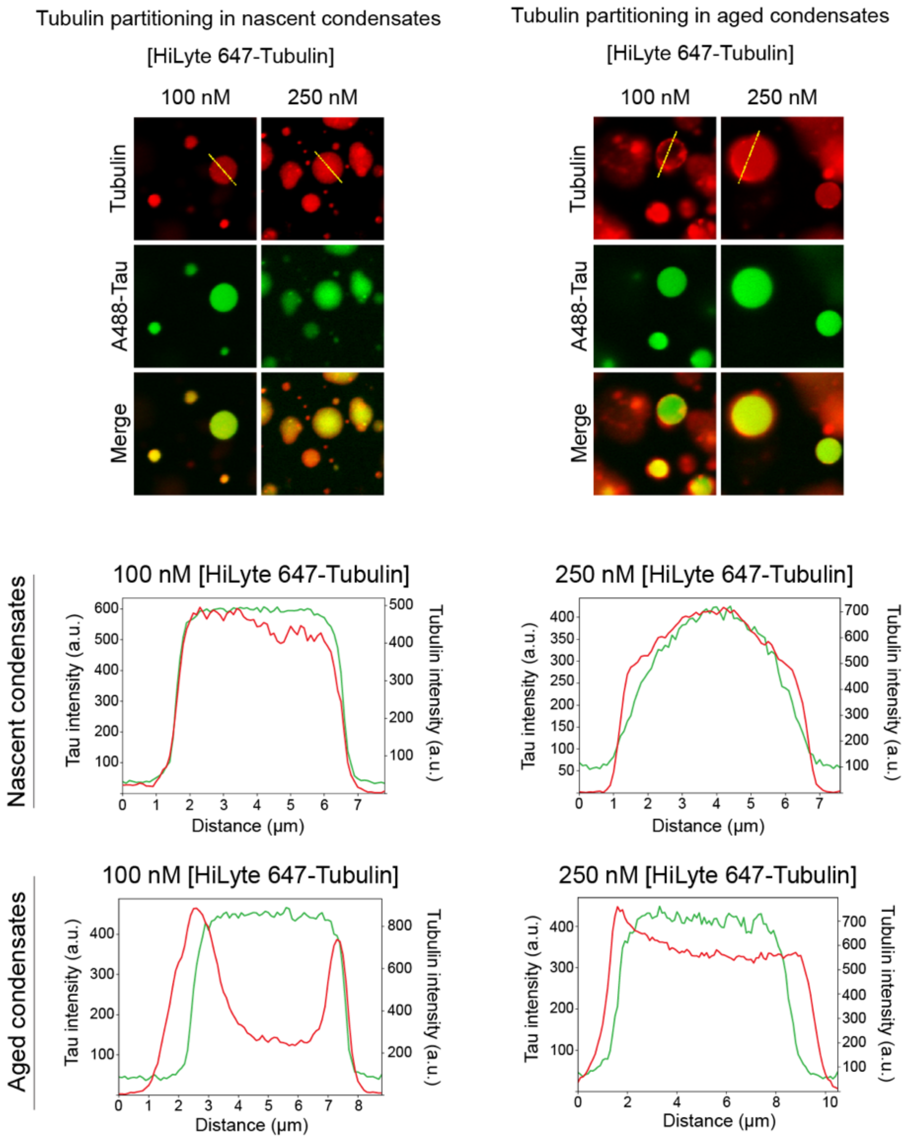


**Supplementary Figure 8**. Partitioning of tubulin (HiLyte647-labeled) in nascent (1 hour since sample preparation) versus aged (4 hours since sample preparation) SynTag-Tau condensates. The concentration of SynTag-Tau protein used is 24 µM with a 10% PEG8000 in a buffer containing 10 mM HEPES (pH 7.4), 50 mM NaCl, 0.1 mM EDTA, and 2 mM DTT along with the specified concentration of HiLyte647-labeled tubulin. Tubulin was added at specific time points after condensate preparation as noted in the figure. The concentration of Atto488-labeled SynTag-Tau is 250 nM. Each experiment was independently repeated three times.

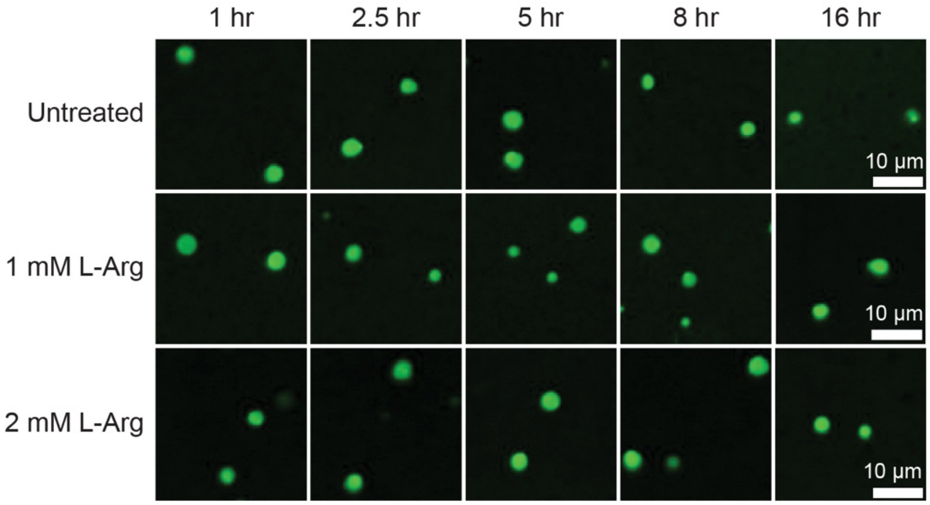


**Supplementary Figure 9**. SynTag-Tau condensates imaged at various time points after sample preparation prior to the addition of tubulin and GTP in MT polymerization assays. The composition of the sample is 12 μM SynTag-Tau in a buffer containing 80 mM PIPES (pH 6.9), 2 mM $MgCl_2$, 0.5 mM EGTA, and 2 mM DTT with 5% PEG 8000. The concentration of Atto 488 labeled SynTag-Tau is 250 nM. This experiment was independently repeated three times.

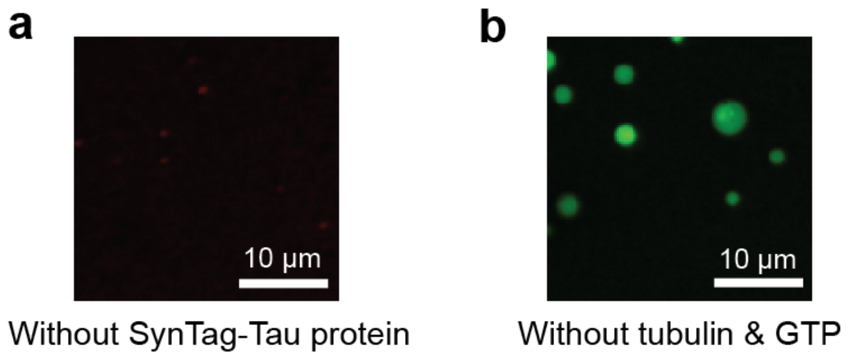


**Supplementary Figure 10**. (a) Lack of visible microtubules (MTs) in the absence of SynTag-Tau condensates. The sample was imaged using HiLyte647-labeled tubulin. It contains 5 µM tubulin and 1 mM GTP in a buffer containing 80 mM PIPES (pH 6.9), 2 mM $MgCl_2$, 0.5 mM EGTA, and 2 mM DTT with 5% PEG 8000. (b) SynTag-Tau condensates, as visualized using Atto488-labeled SynTag-Tau, in the absence of tubulin and GTP (1 hour since sample preparation) in a buffer containing 80 mM PIPES (pH 6.9), 2 mM $MgCl_2$, 0.5 mM EGTA, and 2 mM DTT with 5% PEG 8000. Wherever applicable, the concentrations of Atto488-labeled SynTag-Tau and HiLyte647-labeled tubulin are 250 nM and 500 nM, respectively. Each experiment was independently repeated three times.

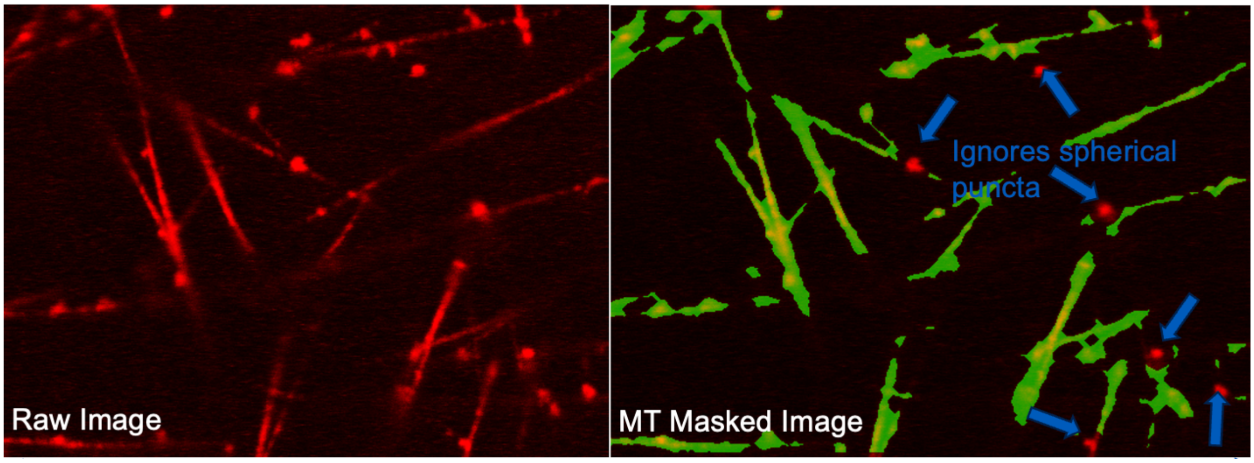


**Supplementary Figure 11**. (left) A fluorescence image of HiLyte647-labeled microtubule (MT) structures formed in the presence of SynTag-Tau condensates. (right) Image analysis allows selective segmentation of filamentous microtubules while ignoring spherical puncta, yielding an estimation of MT surface coverage area. Details of the image analysis protocol are described in the methods section. The composition of the sample on the left panel is 12 μM SynTag-Tau, 5 μM tubulin, and 1 mM GTP in a buffer containing 80 mM PIPES (pH 6.9), 2 mM $MgCl_2$, 0.5 mM EGTA, and 2 mM DTT with 5% PEG 8000. The concentration of HiLyte647-labeled tubulin is 500 nM.

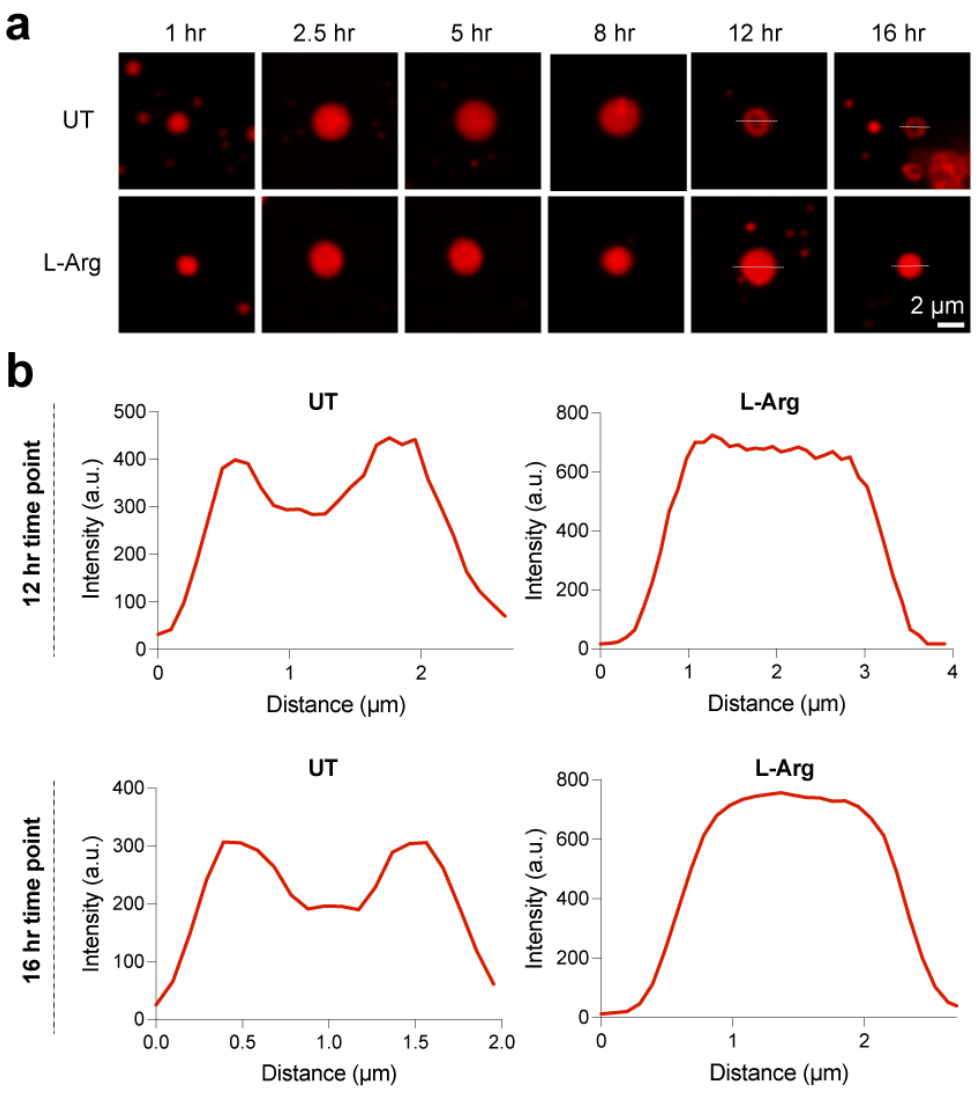


**Supplementary Figure 12.** (a) SynTag-Tau condensates, either untreated (UT) or L-Arg treated (1 mM), imaged at various time points after sample preparation, after the addition of 50 nM HiLyte647-labeled tubulin but not GTP. (b) shows the line profiles corresponding to 12 hr and 16 hr time points, as indicated in (a). The composition of the sample is 12 µM SynTag-Tau in a buffer containing 80 mM PIPES (pH 6.9), 2 mM $MgCl_2$, 0.5 mM EGTA, and 2 mM DTT with 5% PEG 8000, either with or without 1 mM L-Arg. This experiment was independently repeated three times.

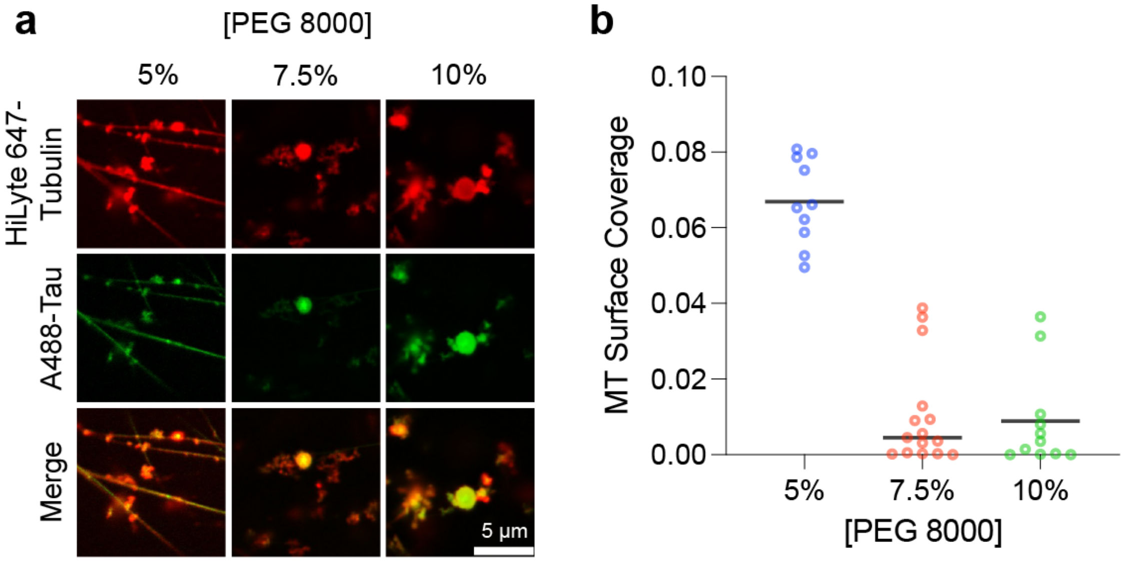


**Supplementary Figure 13**. (a) Representative fluorescence images of microtubule (MT) formation in the presence of SynTag-Tau condensates. These condensates were prepared at variable concentrations of PEG8000 (w/v), as indicated. (b) Corresponding microtubule surface coverage plot as a function of PEG8000 concentration (w/v). The center line represents the median. The individual data points based on 15-17 measurements from three independent replicates are shown. The composition of the sample is 12 μM SynTag-Tau, 5 μM tubulin, and 1 mM GTP in a buffer containing 80 mM PIPES (pH 6.9), 2 mM $MgCl_2$, 0.5 mM EGTA, and 2 mM DTT, along with the indicated concentrations of PEG8000. The concentrations of Atto488-labeled SynTag-Tau and HiLyte647-labeled tubulin are 250 nM and 500 nM, respectively. The sample age in these experiments is 1 hour.

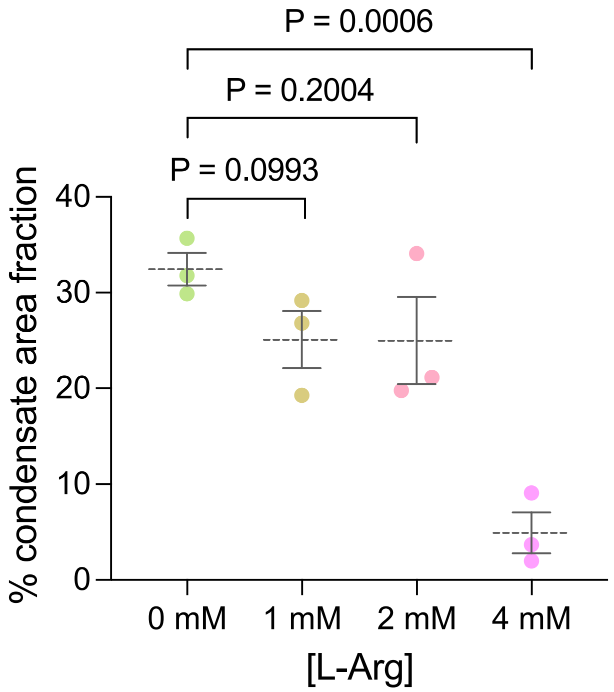


**Supplementary Figure 14.** Estimated SynTag-Tau condensate area fraction (expressed in percentages with respect to the total image acquisition area) at different concentrations of L-Arg, measured at 2 hours after sample preparation. The composition of samples used for this analysis is 12 µM SynTag-Tau in buffer containing 10 mM HEPES (pH 7.4), 50 mM NaCl, 0.1 mM EDTA, and 2 mM DTT, along with 7.5% PEG8000, as well as varied concentrations of L-Arg. The dashed line represents the mean, and the error bars represent the standard error of the mean (S.E.M.) based on three independent replicates, which are represented as individual points. Statistical significance was determined using the unpaired two-sided Student's *t*-test. The respective P values are indicated in the plot.

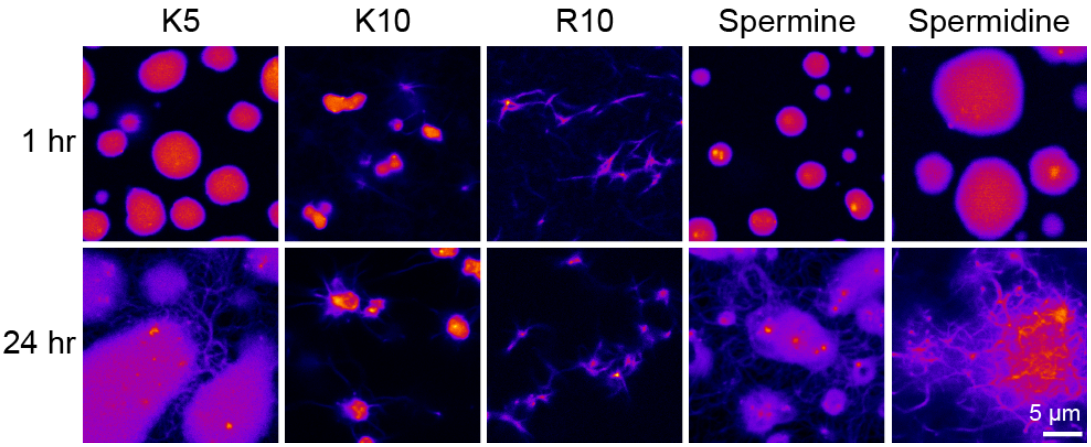


**Supplementary Figure 15**. Effect of multivalent cationic molecules on SynTag-Tau condensate formation and transition to amyloid fibrils. Fluorescence images of these samples were acquired using Atto488-labeled SynTag-Tau. Abbreviations shown in the figure refer to K5 as penta-lysine, K10 as deca-lysine, and R10 as deca-arginine. The composition of the sample is 12 µM SynTag-Tau in buffer containing 10 mM HEPES (pH 7.4), 50 mM NaCl, 0.1 mM EDTA, and 2 mM DTT, along with 7.5% PEG8000 crowder as well as the indicated multivalent molecule at 325 µg/ml (approximately equivalent to 2 mM L-Arg concentration). The concentration of Atto488-labeled SynTag-Tau is 250 nM. Each experiment was independently repeated three times.

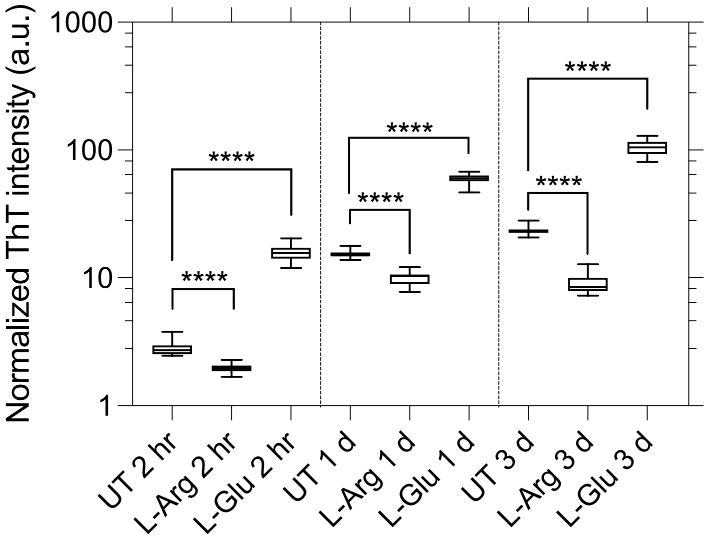


**Supplementary Figure 16**. Wild-type Tau condensate samples with heparin (UT: untreated condition) show an increased Thioflavin T (ThT) fluorescence signal upon aging, which is diminished in the presence of 2 mM L-Arg (L-Arg) and is enhanced in the presence of 2 mM L-Glu (L-Glutamic acid). The time points are indicated in the figure (hr- hours; d- days). The center line in the box plot represents the median based on 15 measurements from three independent replicates, and the whiskers represent the min to max. Statistical significance was determined by performing an unpaired two-sided Student's t-test (**** means $p<0.0001$) between the ThT fluorescence intensities of either UT and L-Arg-treated condensates or UT and L-Glu-treated condensates at different time points. The composition of the sample is 25 μM Tau in buffer containing 10 mM HEPES (pH 7.5), 50 mM NaCl, 0.1 mM EDTA, 2 mM DTT, 7.5% PEG 8000, 6.25 μM heparin, as well as the indicated small molecule at 2 mM (whenever applicable).

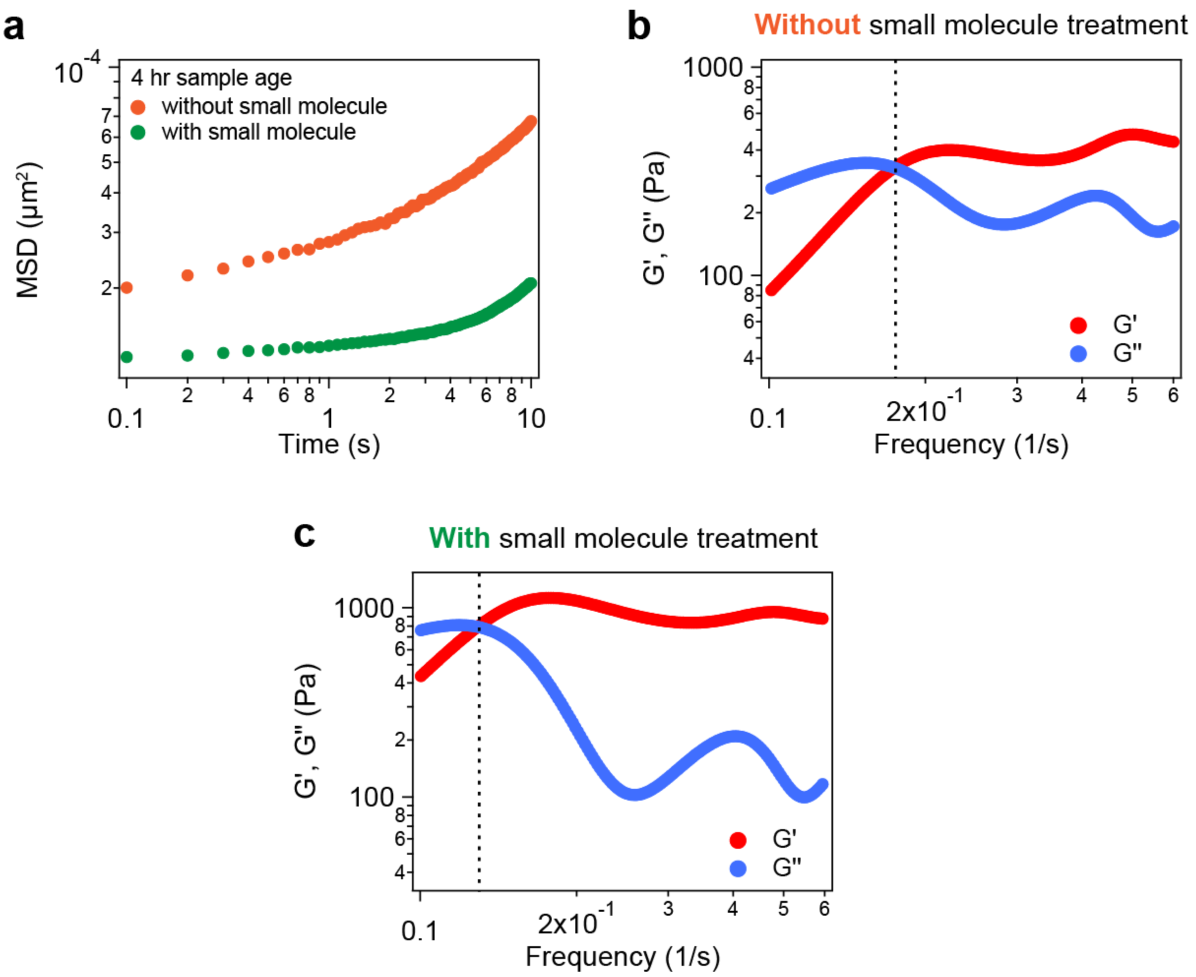


**Supplementary Figure 17**. (a) Representative mean squared displacements (MSDs) of probe particles inside aged Tau condensates, either with or without small molecule treatment (2 mM L-Arg). Representative dynamical moduli of Tau condensates in the absence (b) or presence (c) of 2 mM L-Arg. These measurements were conducted 4 hours after sample preparation. The dashed line in (b) and (c) represents the crossover frequency. The composition of the samples used here is 40 μM Tau in buffer containing 10 mM HEPES (pH 7.4), 50 mM NaCl, 0.1 mM EDTA, and 2 mM DTT along with 7.5% PEG 8000 crowder, and 6.25 μM heparin. Wherever applicable, the L-Arg concentration used in these experiments is 2 mM. The sample size for all these measurements is 3-5 condensates from three independent replicates.

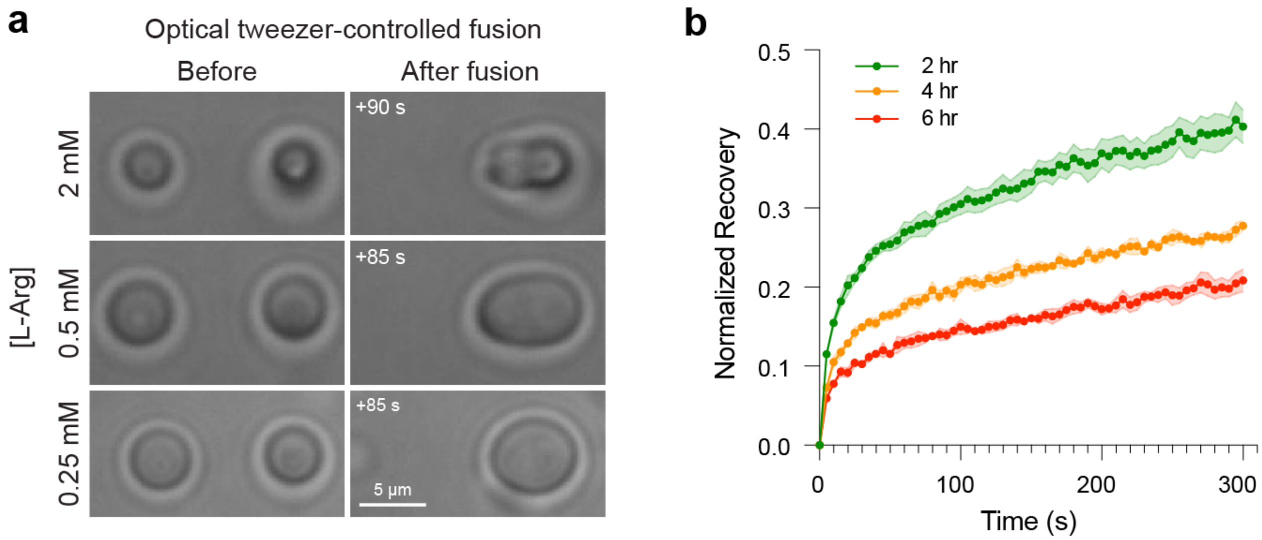


**Supplementary Figure 18**. (a) Optical tweezer-mediated fusion of SynTag-Tau condensates treated with L-Arg at the specified concentrations as indicated. These experiments were conducted 30 minutes after condensate sample preparation. (b) FRAP measurements of 2 mM L-Arg-treated SynTag-Tau condensates, with 250 nM Atto488-labeled SynTag-Tau, at various points of aging (since sample preparation). Intensity error bars were plotted based on the standard error of the mean (S.E.M.) at each time point recorded and represented as the shaded regions. The composition of the samples is 12 µM SynTag-Tau in buffer containing 10 mM HEPES (pH 7.4), 50 mM NaCl, 0.1 mM EDTA, and 2 mM DTT along with 7.5% PEG 8000 crowder. The measurements reported in (a) were repeated independently two times, and those in (b) were repeated independently three times.

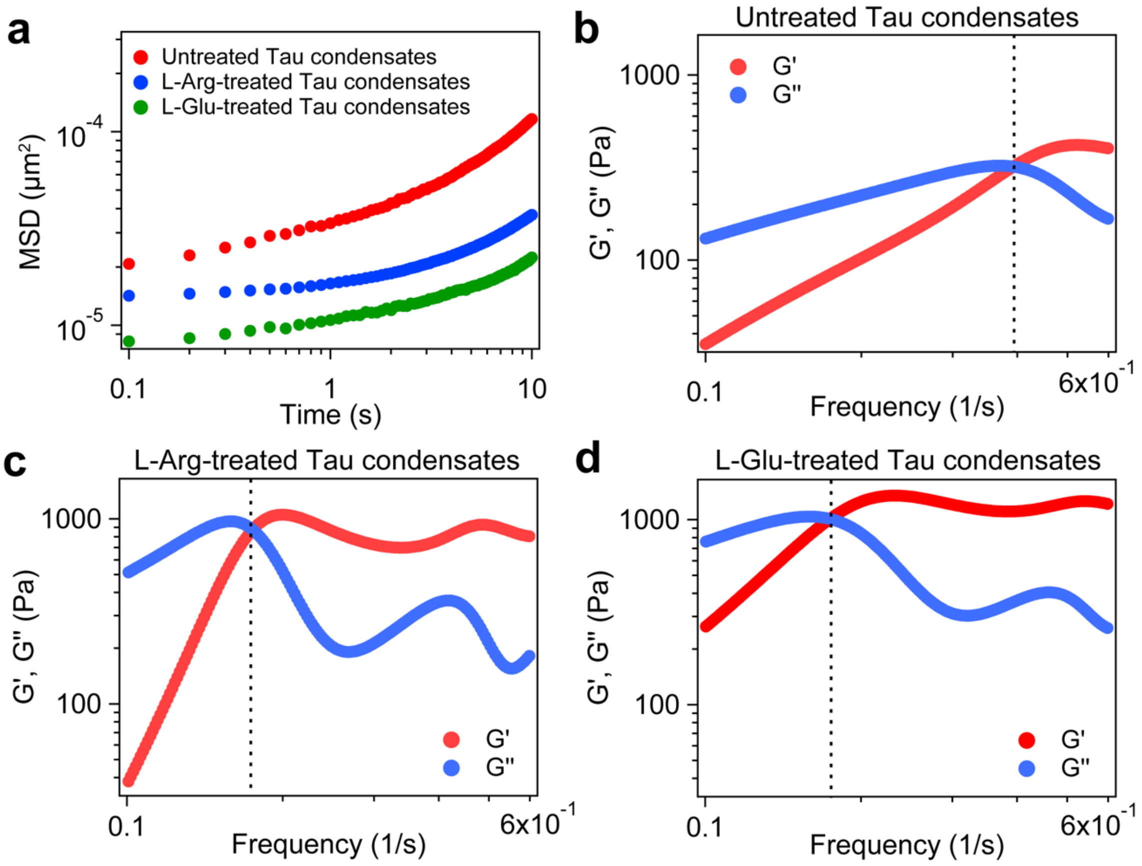


**Supplementary Figure 19.** (a) Representative MSDs of probe particles inside Tau condensates with small molecule treatment [L-arginine, L-Arg or L-glutamic acid, L-Glu] or without small molecule treatment (untreated). Representative dynamical moduli of Tau condensates in the absence (b) or presence of L-Arg (c) or L-Glu (d). These measurements were conducted 2 hours after sample preparation. The 'untreated' and 'L-Arg' data replotted here are based on data reproduced from Fig. 5b, c. The dashed line in (b-d) represents the crossover frequency. The composition of the samples used here is 40 μM Tau in buffer containing 10 mM HEPES (pH 7.4), 50 mM NaCl, 0.1 mM EDTA, and 2 mM DTT, along with 7.5% PEG 8000 crowder, and 6.25 μM heparin. Wherever applicable, the small molecule concentration used in these experiments is 2 mM. The sample size for all these measurements is 3-5 condensates from three independent replicates.

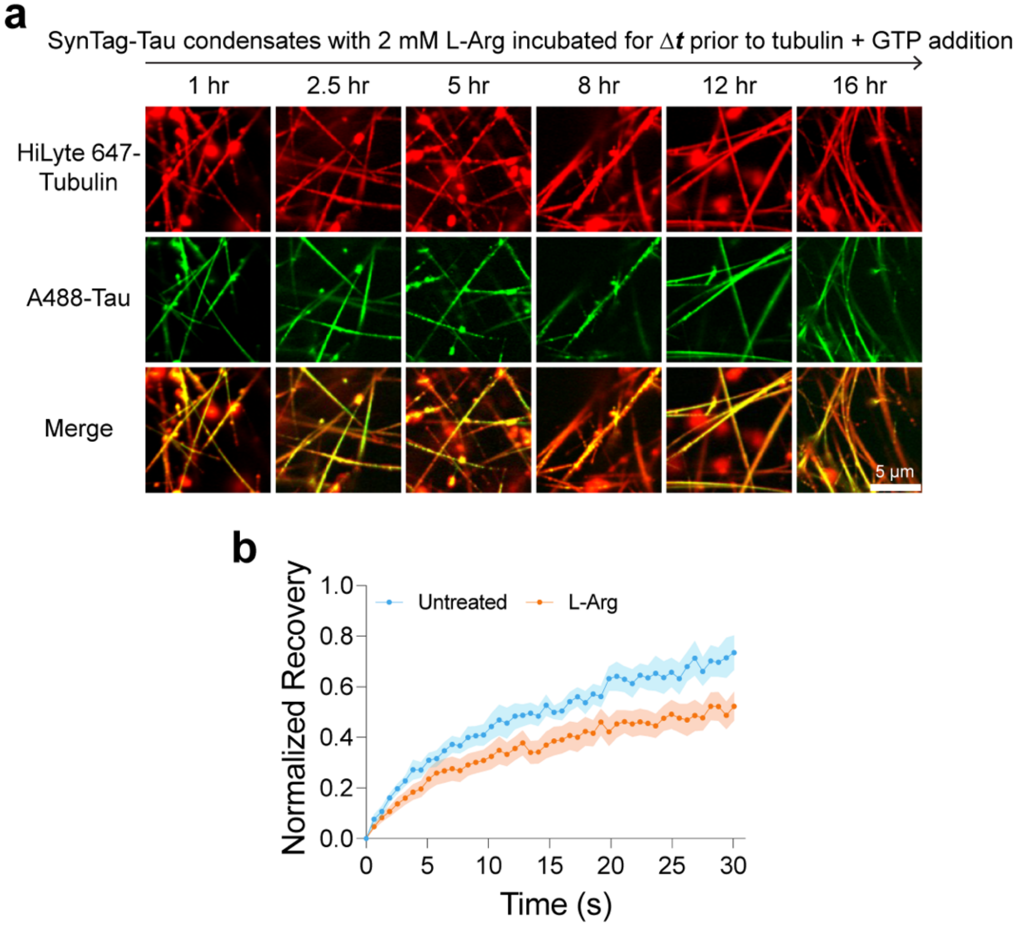


**Supplementary Figure 20.** (a) Microtubule polymerization assay shows that SynTag-Tau condensates treated with 2 mM L-Arg remain active in catalyzing microtubule assembly as compared to untreated condensates after 16 hours of aging. The corresponding quantification is shown in Fig. 6b. The composition of samples used here is: 12 µM SynTag-Tau protein in buffer containing 80 mM PIPES (pH 6.9), 2 mM $MgCl_2$, 0.5 mM EGTA, and 2 mM DTT, along with 5% PEG8000. The concentration of tubulin is 5 µM, and the concentration of GTP is 1 mM. The concentrations of Atto488-labeled SynTag-Tau and HiLyte647-labeled tubulin are 250 nM and 500 nM, respectively. (b) FRAP profiles of HiLyte647-labeled tubulin in SynTag-Tau condensates either without L-Arg (untreated) or with L-Arg (1 mM). Measurements were taken 1 hour after sample preparation. Intensity error bars were plotted based on the standard error of the mean (S.E.M.) at each interval and are represented as the shaded regions. Each of these experiments was independently repeated three times.

### Supplementary Tables

**Supplementary Table 1:** List of proteins and corresponding sequences used in the study.

| S.no | Protein | Sequence |
|---|---|---|
| 1 | Tau | *SNI*MAEPRQEFEVMEDHAGTYGLGDRKDQGGYTMHQDQEGDTDAGLKESPL<br>QTPTEDGSEEPGSETSDAKSTPTAEDVTAPLVDEGAPGKQAAAQPHTEIPE<br>GTTAEEAGIGDTPSLEDEAAGHVTQARMVSKSKDGTGSDDKKAKGADGKTK<br>IATPRGAAPPGQKGQANATRIPAKTPPAPKTPPSSGEPPKSGDRSGYSSPG<br>SPGTPGSRSRTPSLPTPPTREPKKVAVVRTPPKSPSSAKSRLQTAPVPMPD<br>LKNVKSKIGSTENLKHQPGGGKVQIINKKLDLSNVQSKCGSKDNIKHVPGG<br>GSVQIVYKPVDLSKVTSKCGSLGNIHHKPGGGQVEVKSEKLDFKDRVQSKI<br>GSLDNITHVPGGGNKKIETHKLTFRENAKAKTDHGAEIVYKSPVVSGDTSP<br>RHLSNVSSTGSIDMVDSPQLATLADEVSASLAKQGL |
| 2 | SynTag-Tau | MKSSHHHHHHGSSNSSSNNNNNNNNNNNLGIEENLYFQSNIMAEPRQEFEVM<br>EDHAGTYGLGDRKDQGGYTMHQDQEGDTDAGLKESPLQTPTEDGSEEPGSE<br>TSDAKSTPTAEDVTAPLVDEGAPGKQAAAQPHTEIPEGTTAEEAGIGDTPS<br>LEDEAAGHVTQARMVSKSKDGTGSDDKKAKGADGKTKIATPRGAAPPGQKG<br>QANATRIPAKTPPAPKTPPSSGEPPKSGDRSGYSSPGSPGTPGSRSRTPSL<br>PTPPTREPKKVAVVRTPPKSPSSAKSRLQTAPVPMPDLKNVKSKIGSTENL<br>KHQPGGGKVQIINKKLDLSNVQSKCGSKDNIKHVPGGGSVQIVYKPVDLSK<br>VTSKCGSLGNIHHKPGGGQVEVKSEKLDFKDRVQSKIGSLDNITHVPGGGN<br>KKIETHKLTFRENAKAKTDHGAEIVYKSPVVSGDTSPRHLSNVSSTGSIDM<br>VDSPQLATLADEVSASLAKQGL |
| 3 | Shuffled SynTag-Tau | MKSSHHHHHHENLYFQSGSSNSSSNNNNNNNNNNNLGIENIMAEPRQEFEVM<br>EDHAGTYGLGDRKDQGGYTMHQDQEGDTDAGLKESPLQTPTEDGSEEPGSE<br>TSDAKSTPTAEDVTAPLVDEGAPGKQAAAQPHTEIPEGTTAEEAGIGDTPS<br>LEDEAAGHVTQARMVSKSKDGTGSDDKKAKGADGKTKIATPRGAAPPGQKG<br>QANATRIPAKTPPAPKTPPSSGEPPKSGDRSGYSSPGSPGTPGSRSRTPSL<br>PTPPTREPKKVAVVRTPPKSPSSAKSRLQTAPVPMPDLKNVKSKIGSTENL<br>KHQPGGGKVQIINKKLDLSNVQSKCGSKDNIKHVPGGGSVQIVYKPVDLSK<br>VTSKCGSLGNIHHKPGGGQVEVKSEKLDFKDRVQSKIGSLDNITHVPGGGN<br>KKIETHKLTFRENAKAKTDHGAEIVYKSPVVSGDTSPRHLSNVSSTGSIDM<br>VDSPQLATLADEVSASLAKQGL |
| 4 | Truncated SynTag-Tau | *S*GSSNSSSNNNNNNNNNNNLGIENIMAEPRQEFEVMEDHAGTYGLGDRKDQG<br>GYTMHQDQEGDTDAGLKESPLQTPTEDGSEEPGSETSDAKSTPTAEDVTAP<br>LVDEGAPGKQAAAQPHTEIPEGTTAEEAGIGDTPSLEDEAAGHVTQARMVS<br>KSKDGTGSDDKKAKGADGKTKIATPRGAAPPGQKGQANATRIPAKTPPAPK<br>TPPSSGEPPKSGDRSGYSSPGSPGTPGSRSRTPSLPTPPTREPKKVAVVRT<br>PPKSPSSAKSRLQTAPVPMPDLKNVKSKIGSTENLKHQPGGGKVQIINKKL<br>DLSNVQSKCGSKDNIKHVPGGGSVQIVYKPVDLSKVTSKCGSLGNIHHKPG<br>GGQVEVKSEKLDFKDRVQSKIGSLDNITHVPGGGNKKIETHKLTFRENAKA<br>KTDHGAEIVYKSPVVSGDTSPRHLSNVSSTGSIDMVDSPQLATLADEVSAS<br>LAKQGL |

*TEV protease cleavage of the His-tag leaves a few amino acids at the N-terminus of the protein, which is italicized.

*Synthetic prion-like tag sequence is underlined.

**Supplementary Table 2:** Tabulated amide I Lorentzian curve-fitting parameters with peak assignments for BCARS data processing.

| Peak Number | Assignment | Fit Parameters | Seed Value |
|---|---|---|---|
| 0 | Tyrosine Ring Mode | $X_0$ | 1600 ± 5 |
| | | FWHM | 30 |
| | | Area | 1.0 |
| 1 | Tyrosine Ring Mode | $X_0$ | 1612 ± 5 |
| | | FWHM | 30 |
| | | Area | 1.0 |
| 2 | α-helix | $X_0$ | 1644 ± 2 |
| | | FWHM | 24 |
| | | Area | 1.0 |
| 3 | Random Coil | $X_0$ | 1660 ± 2 |
| | | FWHM | 24 |
| | | Area | 1.0 |
| 4 | β-sheet | $X_0$ | 1673 ± 2 |
| | | FWHM | 24 |
| | | Area | 1.0 |
| 5 | β-turn | $X_0$ | 1691 ± 5 |
| | | FWHM | 20 |
| | | Area | 1.0 |

*FWHM is fixed to seed value, and area seed values are constrained to a zero minimum.

### Supplementary Video Legend

**Supplementary Video 1**. The time-dependent transition of SynTag-Tau condensates to fibrils as visualized by Atto488-labeled SynTag-Tau. The concentration of SynTag-Tau used is 24 µM with 10% PEG8000. The buffer composition is 10 mM HEPES (pH 7.4), 50 mM NaCl, 0.1 mM EDTA, and 2 mM DTT. The concentration of Atto488-labeled SynTag-Tau is 250 nM.

### Supplementary References


1. Huang Y, Wen J, Ramirez L-M, Gümüşdil E, Pokhrel P, Man VH, *et al.* Methylene blue accelerates liquid-to-gel transition of tau condensates impacting tau function and pathology. *Nature Communications* 2023, **14**(1)**:** 5444.

2. Schindelin J, Arganda-Carreras I, Frise E, Kaynig V, Longair M, Pietzsch T*, et al.* Fiji: an open-source platform for biological-image analysis. *Nature Methods* 2012, **9**(7)**:** 676-682.

3. Alshareedah I, Kaur T, Banerjee PR. Chapter Six - Methods for characterizing the material properties of biomolecular condensates. In: Keating CD (ed). *Methods in Enzymology*, vol. 646. Academic Press, 2021, pp 143-183.

4. Alshareedah I, Thurston GM, Banerjee PR. Quantifying viscosity and surface tension of multicomponent protein-nucleic acid condensates. *Biophys J* 2021, **120**(7)**:** 1161-1169.

5. Ghosh A, Kota D, Zhou HX. Determining Thermodynamic and Material Properties of Biomolecular Condensates by Confocal Microscopy and Optical Tweezers. *Methods Mol Biol* 2023, **2563:** 237-260.

6. Parekh SH, Lee YJ, Aamer KA, Cicerone MT. Label-free cellular imaging by broadband coherent anti-Stokes Raman scattering microscopy. *Biophys J* 2010, **99**(8)**:** 2695-2704.

7. Day JP, Rago G, Domke KF, Velikov KP, Bonn M. Label-free imaging of lipophilic bioactive molecules during lipid digestion by multiplex coherent anti-Stokes Raman scattering microspectroscopy. *J Am Chem Soc* 2010, **132**(24)**:** 8433-8439.

8. Movasaghi Z, Rehman S, Rehman IU. Raman Spectroscopy of Biological Tissues. *Applied Spectroscopy Reviews* 2007, **42**(5)**:** 493-541.

9. Fleissner F, Bonn M, Parekh SH. Microscale spatial heterogeneity of protein structural transitions in fibrin matrices. *Sci Adv* 2016, **2**(7)**:** e1501778.

10. Chatterjee S, Kan Y, Brzezinski M, Koynov K, Regy RM, Murthy AC*, et al.* Reversible Kinetic Trapping of FUS Biomolecular Condensates. *Advanced Science* 2022, **9**(4)**:** 2104247.

11. Berjot M, Marx J, Alix AJP. Determination of the secondary structure of proteins from the Raman amide I band: The reference intensity profiles method. *Journal of Raman Spectroscopy* 1987, **18**(4)**:** 289-300.

12. Hernández-Vega A, Braun M, Scharrel L, Jahnel M, Wegmann S, Hyman BT*, et al.* Local Nucleation of Microtubule Bundles through Tubulin Concentration into a Condensed Tau Phase. *Cell Rep* 2017, **20**(10)**:** 2304–2312.

13. Savastano A, Flores D, Kadavath H, Biernat J, Mandelkow E, Zweckstetter M. Disease-Associated Tau Phosphorylation Hinders Tubulin Assembly within Tau Condensates. *Angew Chem Int Ed* 2021, **60**(2)**:** 726–730.

14. Hochmair J, Exner C, Franck M, Dominguez-Baquero A, Diez L, Brognaro H*, et al.* Molecular crowding and RNA synergize to promote phase separation, microtubule interaction, and seeding of Tau condensates. *EMBO J* 2022**:** e108882.

15. Quan MD, Liao SJ, Ferreon JC, Ferreon ACM. Fluorescence Lifetime Imaging Microscopy of Biomolecular Condensates. *Methods Mol Biol* 2023, **2563:** 135-148.

16. Mahendran TS, Wadsworth GM, Singh A, Gupta R, Banerjee PR. Homotypic RNA clustering accompanies a liquid-to-solid transition inside the core of multi-component biomolecular condensates. *Nature Chemistry* 2025, **17**(8)**:** 1236-1246.

17. Tinevez J-Y, Perry N, Schindelin J, Hoopes GM, Reynolds GD, Laplantine E*, et al.* TrackMate: An open and extensible platform for single-particle tracking. *Methods (San Diego, Calif)* 2017, **115:** 80-90.

18. Alshareedah I, Moosa MM, Pham M, Potoyan DA, Banerjee PR. Programmable viscoelasticity in protein-RNA condensates with disordered sticker-spacer polypeptides. *Nat Commun* 2021, **12**(6620)**:** 1–14.

19. Alshareedah I, Borcherds WM, Cohen SR, Singh A, Posey AE, Farag M*, et al.* Sequence-specific interactions determine viscoelasticity and ageing dynamics of protein condensates. *Nature Physics* 2024, **20**(9)**:** 1482-1491.

20. Evans R, Tassieri M, Auhl D, Waigh TA. Direct conversion of rheological compliance measurements into storage and loss moduli. *Physical Review E* 2009, **80**(1)**:** 012501.

21. Meng EC, Goddard TD, Pettersen EF, Couch GS, Pearson ZJ, Morris JH*, et al.* UCSF ChimeraX: Tools for structure building and analysis. *Protein Science* 2023, **32**(11)**:** e4792.

22. Abramson J, Adler J, Dunger J, Evans R, Green T, Pritzel A*, et al.* Accurate structure prediction of biomolecular interactions with AlphaFold 3. *Nature* 2024, **630**(8016)**:** 493-500.

23. Goldschmidt L, Teng PK, Riek R, Eisenberg D. Identifying the amylome, proteins capable of forming amyloid-like fibrils. *Proceedings of the National Academy of Sciences* 2010, **107**(8)**:** 3487-3492.

24. Knox C, Wilson M, Klinger CM, Franklin M, Oler E, Wilson A*, et al.* DrugBank 6.0: the DrugBank Knowledgebase for 2024. *Nucleic Acids Res* 2024, **52**(D1)**:** D1265-d1275.

25. Lancaster AK, Nutter-Upham A, Lindquist S, King OD. PLAAC: a web and command-line application to identify proteins with prion-like amino acid composition. *Bioinformatics* 2014, **30**(17)**:** 2501-2502.
